# Synaptic changes contribute to persistent extra-motor behaviour deficits in the rNLS8 TDP-43 mouse model of amyotrophic lateral sclerosis

**DOI:** 10.1101/2025.03.04.641357

**Authors:** Wei Luan, Rebecca San Gil, Lidia Madrid, Maize C. Cao, Florencia Vassallu, Heledd Brown-Wright, Adekunle T. Bademosi, Yi Jia Chye, Hao Yu Wu, Mu Sheen Chang, Emma L. Scotter, Catherine A. Blizzard, Lionel M. Igaz, Adam K. Walker

## Abstract

Extra-motor symptoms are increasingly recognised in amyotrophic lateral sclerosis (ALS), encompassing cognitive, social, and behavioural deficits that can substantially impact quality of life. TAR DNA binding protein 43 (TDP-43) pathology is the central disease marker of almost all cases of ALS and approximately half of all frontotemporal dementia (FTD). However, the mechanisms linking TDP-43 pathology with extra-motor symptoms in TDP-43-associated neurodegenerative diseases remain unresolved. In this study, we used the rNLS8 mouse model, which expresses human TDP-43 with an ablated nuclear localisation sequence (hTDP-43^ΔNLS^) in a doxycycline-regulatable manner causing progressive motor decline reminiscent of ALS, to delineate the molecular changes associated with disease-relevant phenotypes. We found that in addition to previously reported dramatic motor decline, rNLS8 mice also develop extra-motor phenotypes consistent with FTD, including disinhibition-like and anxiety-like behaviours, and social interaction impairments. These changes began in the earliest disease stages and remained readily detectable even when rNLS8 mice became severely motor impaired. Notably, extra-motor deficits persisted in rNLS8 mice that had recovered motor function upon hTDP-43^ΔNLS^ transgene suppression, regardless of whether recovery was initiated at timepoints prior to or after overt neurodegeneration begins. Transcriptomic analysis of rNLS8 mouse cortex tissues revealed early alterations in expression of 321 genes, most notably involving neuroinflammatory-related pathway activation, and all but 2 of these genes returned to control levels upon suppression of hTDP-43^ΔNLS^ expression. Further, 814 genes showed differential exon usage, indicating changes in alternative splicing, in rNLS8 mouse cortex. Of these, differential exon usage of 10 neuronal genes persisted after hTDP-43^ΔNLS^ transgene suppression, including synapse component genes *Nrxn1*, *Unc13a*, and *Gls*. Similarly, proteomics analysis of the cortex of rNLS8 mice revealed depletion of synaptic proteins, particularly those involved in glutamatergic signalling pathways, which also persisted following hTDP-43^ΔNLS^ transgene suppression. Similar glutamatergic pathway changes were detected in human ALS and FTD post-mortem cortex tissues. Our findings indicate that extra-motor phenotypes emerge early in disease in rNLS8 mice and remain evident despite progressive motor impairments. Further, extra-motor phenotypes persist even upon motor recovery, correlating with specific synaptic gene expression and splicing changes. Overall, this study suggests the potential utility of an expanded suite of behavioural paradigms in preclinical testing using rNLS8 mice, with enhanced relevance to the diversity of TDP-43 proteinopathies including FTD. Our findings further suggest that targeting glutamatergic synaptic components may be an avenue to correct extra-motor deficits associated with TDP-43 pathology.

## Introduction

Amyotrophic lateral sclerosis (ALS) is the most common motor neuron disease (MND), with relentlessly progressive decline due to neuromuscular degeneration, and average ∼3-year lifespan after diagnosis^1^. In addition to the predominant motor manifestation, extra-motor changes, such as disinhibition and alterations in social and executive function, are now recognised as important features of ALS^2^. This appreciation of extra-motor components has resulted in a proposed revision of ALS diagnostic criteria to incorporate neuropsychological deficits^3, 4^. Indeed, up to 65% of people living with ALS develop extra-motor cognitive and behavioural symptoms^5^, and up to 15% of people with ALS also meet the criteria for diagnosis of frontotemporal dementia (FTD)^6^. Notably, extra-motor symptoms adversely impact quality of life, resulting in cognitive, linguistic, mood, and social deficits, and are associated with shorter survival in people living with ALS^7^. Nevertheless, only ∼1% of ALS research has focused on understanding these extra-motor symptoms^8^ and less than 5% of ALS clinical trials have assessed neuropsychological symptoms as outcome measures^9^. Disease-modifying therapies for people living with FTD in the absence of ALS-associated motor impairments are also lacking. This suggests a need for better understanding of the mechanisms driving extra-motor manifestations in these diseases to inform future therapeutic avenues. Likewise, animal models recapitulating extra-motor features in the context of concomitant motor decline are required for preclinical assessment of relevance to the full experience of ALS^10^.

Cytoplasmic mislocalisation and accumulation of TAR DNA-binding protein 43 (TDP-43) have been observed in up to 97% individuals with ALS and nearly 50% FTD cases^11^. Loss of TDP-43 function in the nucleus and accumulation of aggregated TDP-43 in the cytoplasm are concurrent events in neurons that are strongly linked to neurodegeneration^11^. However, the precise mechanisms by which TDP-43 pathology induces disease remain unclear. Evidence suggests that a large number of TDP-43 target genes are related to synaptic organisation and function^12, 13^, disturbances of which appears to contribute to disease development and symptoms in ALS and FTD^14, 15^. Recent findings also highlight TDP-43-mediated RNA splicing and processing of critical axonal and synaptic components associated with neurodegeneration, for example, STMN2^16–18^ and UNC13A^19–22^. It remains to be determined whether changes in levels of synaptic proteins are associated with extra-motor behavioural phenotypes in disease.

The ‘regulatable NLS’ (rNLS8) TDP-43 mouse model, which expresses human TDP-43 with a defective nuclear localisation sequence (NLS), hTDP-43^ΔNLS^, under doxycycline (dox)-suppressible promoter directed to neurofilament heavy chain (NEFH)-positive neurons, has become an established model for understanding the role of TDP-43 pathology in disease^23–27^. rNLS8 mice accumulate cytoplasmic insoluble TDP-43, and develop progressive motor neuron loss, dramatic motor dysfunction and decreased survival, which can be functionally restored by suppression of hTDP-43^ΔNLS^ expression^28^. In the rNLS8 mouse model, synaptic proteins are significantly decreased from 1 week off dox (WOD)^29^, and significant neuronal loss is observed in cortical^28^ and hippocampal^24^ regions from 3-4 WOD, with later loss of spinal cord alpha motor neurons^28^. Notably, extra-motor behavioural deficits develop in transgenic mice with *CamkIIa* promoter brain-restricted hTDP-43^ΔNLS^ expression that lack typical ALS-like motor decline^30^, and rNLS8 mice have been shown to develop features of hyperexcitability, hyperactivity and cognitive deficits^24, 25^. However, despite the known molecular, cellular, and motor deficits that recapitulate ALS in rNLS8 mice, the extra-motor phenotypes of this model have not been fully examined and the molecular mechanisms driving such features of disease are unknown, particularly during functional recovery.

In this study, we characterised extra-motor phenotypes and their molecular drivers in rNLS8 mice after short-term and long-term induction of neuronal hTDP-43^ΔNLS^ expression. Firstly, we observed that short-term expression of hTDP-43^ΔNLS^ led to persistent motor and extra-motor behavioural phenotypes in the rNLS8 mice that unexpectedly remained after motor recovery. Cortex transcriptomics showed that these behavioural phenotypes correlated with significant differential exon usage of neuronal genes, linking the dysregulation of splicing driven by TDP-43 RNA-binding with persistent extra-motor phenotypes. Furthermore, long-term induction of hTDP-43^ΔNLS^ in rNLS8 mice caused persistent disinhibition-like behaviour, executive and habituation deficits, increased anxiety-like behaviour, and social impairment during and even after motor recovery. These behaviours correlated with a significant decrease in glutamatergic synapse components, suggesting that glutamatergic synapse deficits may underly extra-motor phenotypes. Further, the set of glutamatergic synapse genes/proteins identified were also significantly decreased in published transcriptomic and proteomic datasets from human ALS and FTD post-mortem tissues. Overall, our data indicate that rNLS8 mice develop readily detectable extra-motor phenotypes independent of and despite motor decline, and display molecular signatures of TDP-43 loss-of-function related to changes in synaptic protein levels. Importantly, not all behavioural and molecular phenotypes were readily reversible in this model, which is a relevant consideration for maintenance of quality of life for people with disease in a future with therapies that can effectively target TDP-43.

## Methods

### Animals

rNLS8 mice were produced from the intercross of >10 generation back-crossed C56BL/6JAusb background homozygous *tetO*-hTDP-43^ΔNLS^ line 4 mice (B6.C3-Tg(tetO-TARDBP*)4Vle/JAusb, derived from Jackson Laboratories stock #014650) with hemizygous *NEFH*-tTA line 8 mice (B6.C3-Tg(NEFH-tTA)8Vle/JAusb, derived from Jackson stock #025397), constantly fed with dox-containing chow (200mg/kg, Specialty Feeds, Australia)^28, 31^. Experiments were conducted with approval from the Animal Ethics Committee of The University of Queensland (#QBI/131/18 and 2022/AE000578). During experiments, male and female mice were switched to normal chow (SF00-100, Specialty Feeds, Australia) to induce expression of hTDP-43^ΔNLS^ at approximately ten weeks of age. All mice were housed in temperature- and humidity-controlled conditions (21 ± 1 °C, 55 ± 5%) with a 12-h light/dark cycle (lights on at 06:00 h).

### Behaviour tests

#### Neurological score

Mice were assessed for observation of collapsing splay or clasping of hindlimbs on a scale of 0 to 4 by tail suspension for >5 s as previously described^28^.

#### Open field test

Open Field Assessment of general exploratory locomotion in a novel clear Plexiglas (30 cm x 30 cm x 50 cm) arena with white floor divided by two zones: periphery and centre (comprising 50% of the total area cantered) as previously described^30^. The activities were assessed during 30 min with illumination 30 LUX and analysed by time (bin = 5 min).

#### Social interaction test

Social interaction was assessed using a social approach test in a three chamber apparatus with identical Plexiglas chambers (40 × 40 × 40 cm) as previously described^32^. Before testing, the animal was allowed to explore freely for 5 min in the testing arena which served to reduce novelty-induced locomotor hyperactivity. During the sociability testing session, one wire cage contained an unfamiliar C57BL/6JAusb mouse (same sex as test mouse), and the other one contained a novel object made of black LEGO™ (Billund, Denmark). A camera above the maze captured videos. Social interaction time was analysed as nose orientation towards the wire cage within a 2-cm interaction zone adjacent to the wire cage which was outlined EthoVision analyses (Noldus, The Netherlands). During the social recognition session for assessment of social memory, the inanimate object was replaced by a new mouse as the ‘novel’ mouse, whereases, the previous ‘novel’ mouse became the ‘familiar’ mouse. The testing animal was placed in the centre of the middle chamber to explore for 5 minutes. Video records were defined and analysed as above by a trained experimenter who was blinded to the treatment.

#### Y-maze test

A Y maze with three identical arms made of transparent Plexiglas (25 cm × 25 cm × 25 cm) placed at 120° angles to each other was used and placed in a room with clues to allow for visual orientation with illumination 30 lux as previously described^33^. Each mouse was placed at the end of one arm facing the centre and allowed to explore the maze freely for 8 min without training, or reward, while the experimenter remained out of sight. The percentage of spontaneous alternation was defined as the number of actual alternations divided by the possible alternations [(# alternations)/(total arm entries − 2) × 100]. Total entries were scored as an index of ambulatory activity in the Y maze and mice with scores below 12 were excluded as previously outlined^33^.

### RNA extraction and analysis

#### RNA extraction

RNA was extracted with Qiazol (Qiagen #79306) using the Qiagen RNeasy Mini Kit (Qiagen, #74104) and Precellys tissue homogeniser (Bertin Instruments, Montigny-le-Bretonneux, France). On-column DNase I digestion was conducted using RNase-free DNase I (Qiagen #79254). The concentration of extracted RNA was determined using a NanoRNA kit (Agilent #5067-1511) and Bioanalyzer (Agilent 2100, Santa Clara, CS, USA). cDNA was synthesised from 1 μg total RNA using the SuperScript™ VILO™ Master Mix (Thermo Fisher #11755050).

#### Transcriptomic analysis

RNA was purified from the left rostral cortex of control and rNLS8 mice (*n* = 4/group) at disease onset (2 WOD) and recovery (2 WOD followed by 6 weeks on dox). The quality of total RNA samples was determined by Agilent Bioanalyzer with the RNA Integrity Number (RIN) >8.0 and were then used for library construction. RNA was analysed by RNA-seq (polyA enriched, strand-specific, Illumina paired-end 150 bp, 20 M reads) with Azenta Life Science. Data was trimmed (Trimmomatic) for differential expression analysis (DESeq2^34^), and differential exon usage analysis (DEXSeq^34^).

### Meta-analysis of proteomics data

To understand the molecular signatures of the cortex during recovery we re-analysed a published longitudinal proteomics dataset of the rNLS8 cortex^29^. Briefly, a subset of proteins that were significantly decreased in rNLS8 cortex in late disease (6 WOD) were analysed for their protein abundance at recovery (6 WOD followed by 2 weeks on dox). These proteins were then grouped into subsets of proteins that demonstrated “recovered” (i.e., returned to control levels) or “persistent” (i.e., remained significantly decreased) signatures. The relative protein abundance of each protein was presented using the ComplexHeatmap^35^ package in RStudio.

### Gene ontology analysis

Protein or gene subsets were deposited into Metascape ^36^ or QIAGEN Ingenuity Pathway Analysis (version 01-21-03) ^37^ for analysis of enriched biological terms and identification of protein-protein interaction networks.

### Statistical analysis

For two-value data, statistical analyses were conducted using a two-tailed t-test. Two-way ANOVA was used for analyses of datasets with four values with Bonferroni’s post hoc test using Prism-GraphPad software (version 9). Statistical significance is indicated as \**p* < 0.05, \*\**p* < 0.01, and \*\*\**p* < 0.001. Data are presented as mean ± standard error of the mean (SEM) or ± standard deviation (SD) as shown in the figure legends.

### Data availability statement

The transcriptomic data will be deposited to the Gene Expression Omnibus repository (dataset identifier to be updated). Previously published mouse proteomics^29^, human proteomics^38^ and human transcriptomics^39^ datasets used in this work for comparative analyses are available from the respective references. Source data and Supplementary Data are provided with this paper and any additional information required to re-analyze the data reported in this paper is available from the lead contact upon request.

## Results

### Short-term TDP-43 cytoplasmic accumulation induces persistent hyperlocomotion in rNLS8 mice

Synaptic dysfunction is increasingly recognised as a contributor to extra-motor phenotypes preceding neurodegeneration in ALS and FTD^13,14^. Notably, TDP-43 pathology has been implicated in disturbances of synaptic function, neuronal activity, and behavioural alterations^14, 40^. To investigate whether behavioural changes are dependent on neuron loss or are triggered by the contribution of TDP-43 cytoplasmic mislocalisation to extra-motor symptoms prior to detectable neurodegeneration, we examined motor and extra-motor behaviours in “short-term diseased” rNLS8 mice induced to express hTDP-43^ΔNLS^ for only 2 weeks off dox (WOD), a timepoint prior to loss of cortical or spinal cord neurons^28^, followed by 6 weeks back on dox to supress transgene expression (Fig. 1A).

**Figure 1.**
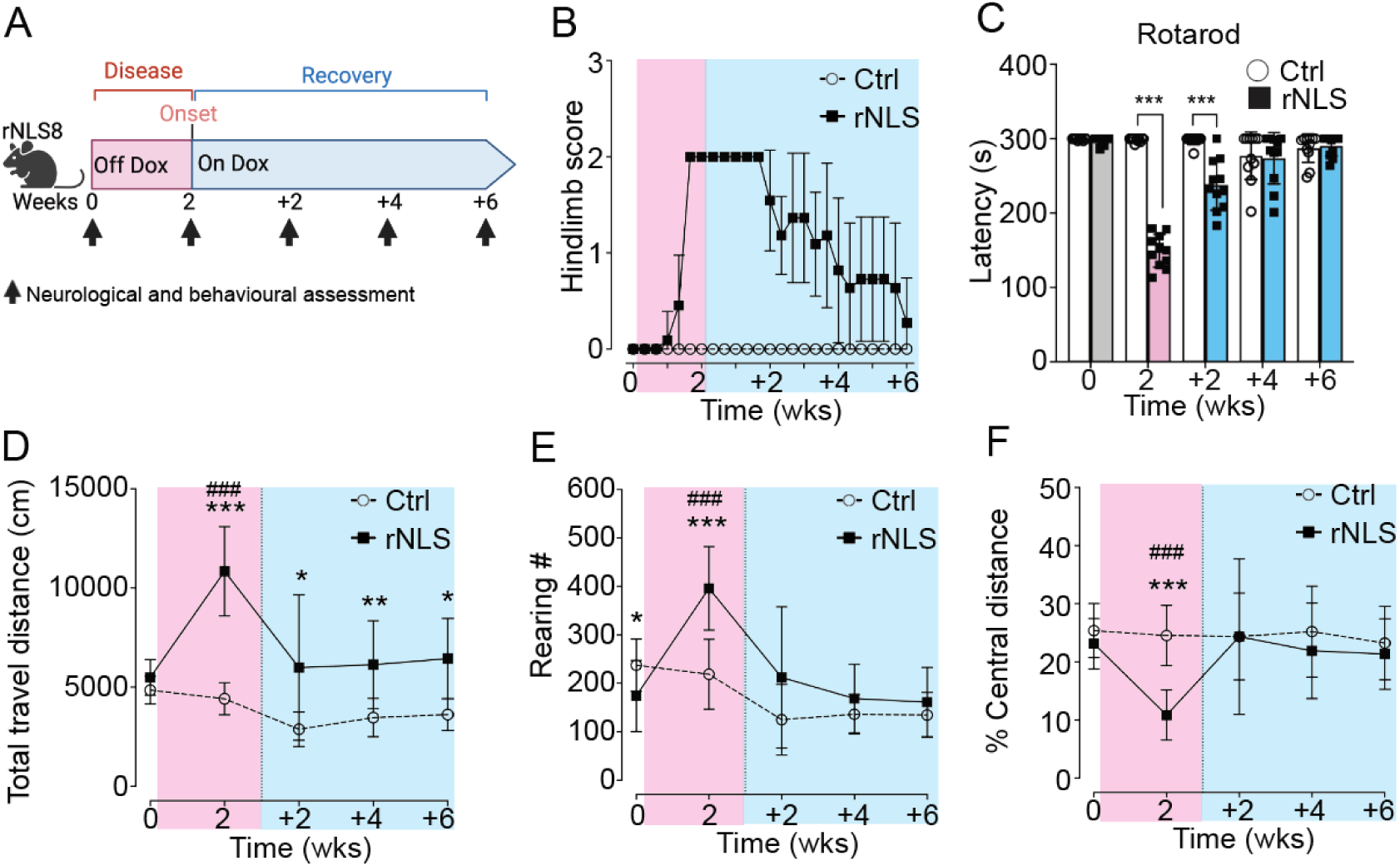
rNLS8 mice at disease onset show recoverable motor deficits but persistent hyperlocomotion. (**A**) Experimental schematic. Mice were behaviourally evaluated at 0 and 2 WOD, and +2, +4, +6 weeks back on dox. Results for the hindlimb neurological score (**B**) and latency to fall in seconds in the rotarod test (**C**) are shown. Mice that explored the open field arena for over 30 min were analysed for the total distance travelled (**D**), rearing number (**E**) and relative central travel distance (**F**). Data are shown as mean ± SD. n = 11. * p < 0.05, * p < 0.01, *** p < 0.001 between control and rNLS8 mice at the same timepoints by repeated one-way ANOVA. ### p < 0.001 of rNLS8 mice relative to other timepoints by repeated one-way ANOVA.

As expected, rNLS8 mice displayed a neurological score of 2 indicating collapsed hindlimb splay by 2 WOD, which recovered at 2 WOD +6 wks (Fig. 1B). Likewise, motor performance on the rotarod test declined at 2 WOD and was fully restored by +4 wks to a level indistinguishable from control mice (Fig. 1C). Overall, the motor behaviour results indicate early motor decline that is readily restored towards control levels over 6 weeks of recovery, similar to previous findings of motor function recovery when rNLS8 mice are returned to dox at later timepoints^28^.

Having established the expected motor decline and recovery phenotypes in rNLS8 mice, we next initially assessed locomotor activity, anxiety-like behaviours, and general exploratory activities using the open field test^41, 42^. At 2 WOD, rNLS8 mice exhibited increased travel distance suggesting hyperlocomotion, and increased rearing frequency and decreased central travel distance, potentially indicating a disinhibition-like phenotype^41^ (Fig. 1D-F). Further, rNLS8 mice remained significantly more hyperactive than controls across the recovery time course, with only partially normalised hyperlocomotion after 6 weeks back on dox (Fig. 1D), indicating enduring hyperlocomotion (Supplementary Fig. 1A). Interestingly, the increased rearing frequency (Fig. 1E) and decreased relative central travel distance (Fig. 1F) were completely restored to control levels by 2WOD + 6 wks (Supplementary Fig. 1B,C). Overall, these findings suggest that the short-term hTDP-43^ΔNLS^ expression was sufficient to induce motor and extra-motor behavioural phenotypes, some of which persisted in the recovery phase.

### Early and persistent differential exon usage in neuronal genes correlates with behavioural abnormalities in rNLS8 mice

To identify molecular changes associated with the enduring hyperlocomotion phenotype in the rNLS8 mice, we conducted bulk RNA sequencing of cortex tissue. Given that TDP-43 acts as an RNA-binding protein regulating both expression and splicing, we assessed changes to gene expression and differential exon usage (DEU; a measure of alternative splicing) in the cortex of mice at disease onset (2 WOD) and in the “short-term disease” recovery experimental paradigm (2 WOD + 6 wks, Fig. 2).

**Figure 2.**
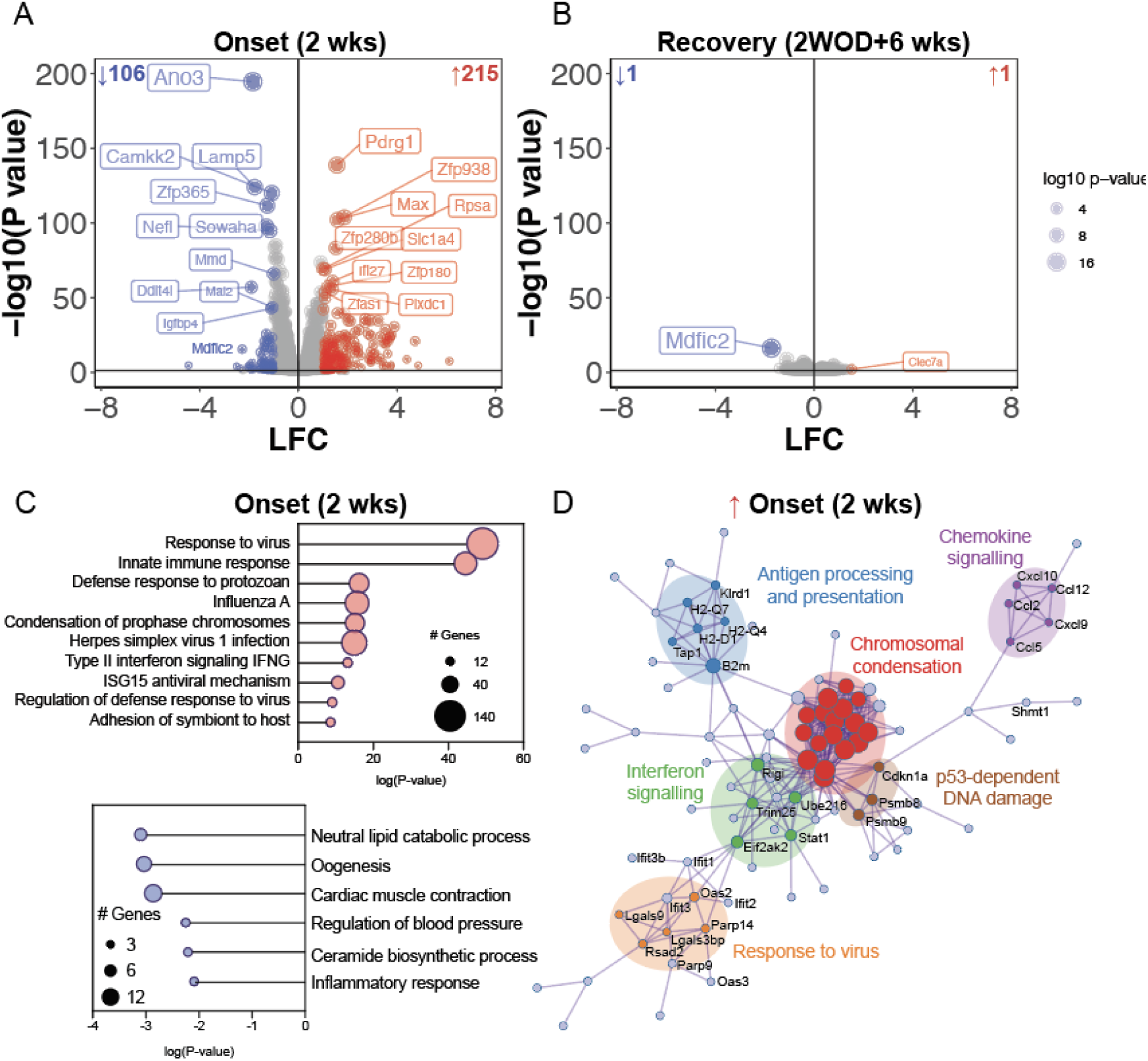
The cortical transcriptome at disease onset is strongly associated with neuroinflammation and this is completely normalised in recovery in rNLS8 mice. (**A-B**) Volcano plots of transcript mean log fold change (LFC; rNLS8/Con) and significance level [-log10(Pvalue)] of cortex from (**A**) disease onset (2 WOD) and (**B**) recovery (2 and 6 WOD). Significantly increased genes are in red and decreased in blue. (**C**) Gene ontology analysis of biological process enrichment at disease onset (2 WOD) of significantly increased (top) and decreased (bottom) genes. Size of the data points are relative to the number of genes in the term. (**D**) Protein-protein interaction networks of significantly increased genes at disease onset (2 WOD). Genes are coloured by membership to given biological processes. Data is from n = 4 mice per group.

At disease onset, *n* = 321 genes were significantly altered in the cortex (log fold change > 1, P-value <0.05) including *n* = 106 decreased and *n* = 215 increased (Fig. 2A, Supplementary Table 1). In contrast, after 6 weeks of recovery there were negligible transcriptomic differences between control and rNLS8 mouse cortex (Fig. 2B). Indeed, only 2 genes were persistently altered in 2 WOD +6 weeks rNLS8 mice, namely increased *Clec7a* (encoding C-type lectin domain containing 7A) and decreased *Mdfic2* (encoding MyoD Family Inhibitor Domain Containing 2) at both 2 WOD and 2 WOD +6 weeks back on Dox. Differentially expressed genes at disease onset (2 WOD) were strongly associated with a decrease in lipid metabolism (lipid catabolic and ceramide biosynthetic processes, Fig. 2C) and an increase in inflammatory pathways (response to virus and innate immune response), with a highly connected protein-protein interaction network (Fig. 2D). The contrast in the number of differentially expressed genes between disease onset (2 WOD) and recovery (2WOD +6 wks) demonstrates the efficient regulation of the dox-suppressible hTDP-43^ΔNLS^ transgene and the capacity of the cortex to recover to the control transcriptome. Nevertheless, there were no differentially expressed genes that persisted from onset to recovery from “short-term disease” that could explain the persistent hyperlocomotion phenotype in the rNLS8 mice.

We then hypothesised that alternative splicing changes due to TDP-43 dysfunction may persist from disease onset to recovery to drive enduring behavioural changes in the rNLS8 mice. We identified *n* = 814 genes with significant differential exon usage (DEU) in the cortex of rNLS8 mice after only 2 weeks of hTDP-43^ΔNLS^ expression (Fig. 3). Genes with DEU events were enriched for cell projection organisation, synaptic signalling (including a significant network of glutamatergic synapse genes), and pathways involved in neurodegeneration such as ER to Golgi anterograde transport, vesicle fusion, neddylation, and regulation of proteolysis (Fig. 3A,B).

**Figure 3.**
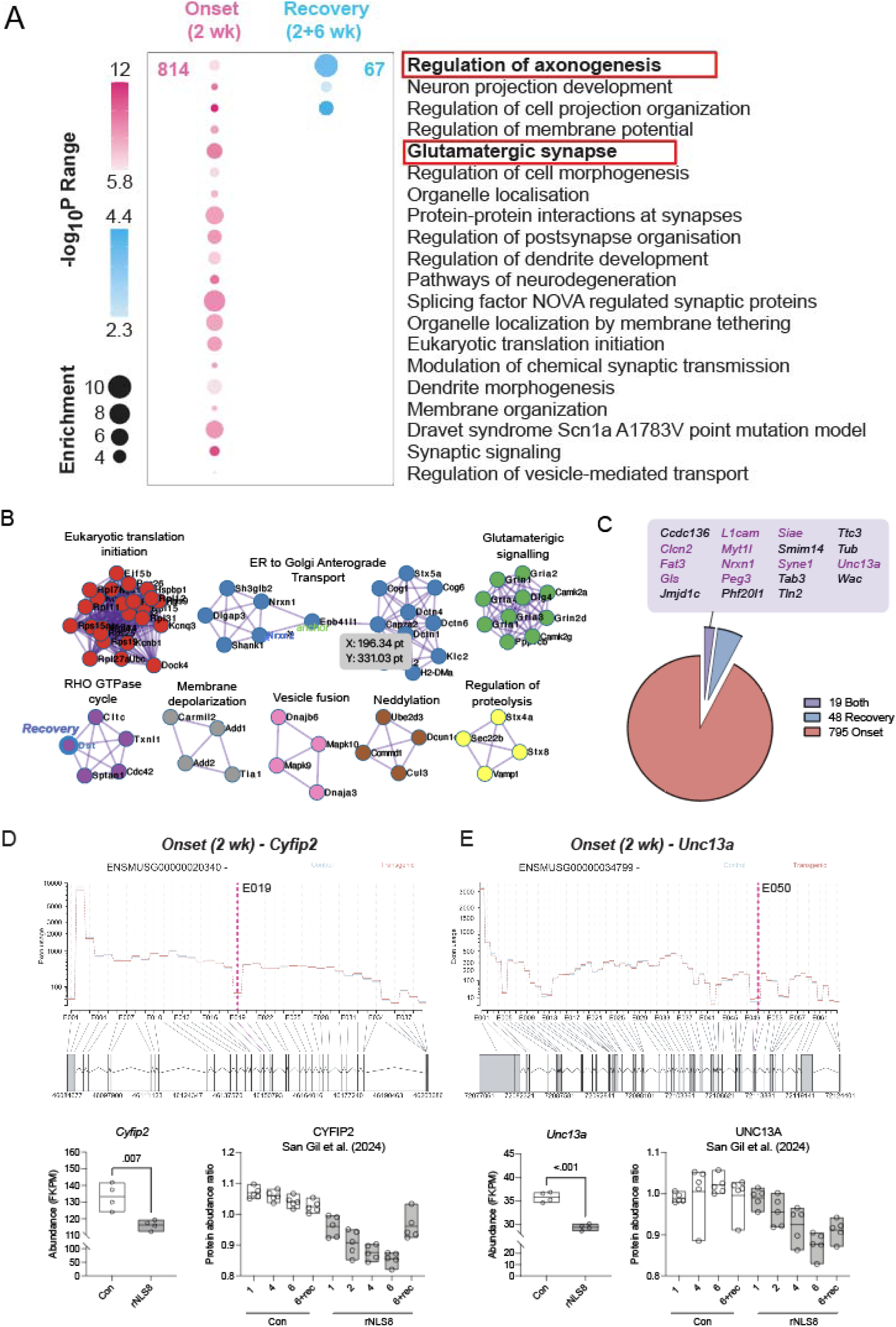
Differential exon usage in genes associated with axonogenesis and neuronal projection organisation at disease onset and in recovery cortex of rNLS8 mice. (**A**) Metascape gene ontology of onset (n = 814) and recovery (n = 67) genes that show signficiant alternative splicing. Bubble sizes increase with enrichment ratio and colour gradients are scaled to the range of P-values in these (pink) and recovery (blue) genes, with darker colours representing stronger significance levels. (**B**) Protein-protein interaction networks of genes with significant DEU events. All genes and networks relate to DEU genes in onset (2 WOD) cortex with the exception of Dst which also has DEU in recovery cortex (outlined in blue, member of the RHO GTPase cycle). (**C**) Proportion of genes with DEU events at onset (red), recovery (blue) or both onset and recovery (purple). List of genes with DEU events in both onset and recovery cortex are listed alphabetically and neuronal genes are in purple. (**D**) Differential exon usage of Cyfip2 at E019 correlates with a significant decrease in Cyfip2 expression (fragments per kilobase of transcript per million; FKPM). Analysis of a published longitudinal proteomic dataset from control and rNLS8 cortex ^29^ shows a gradual decrease in cytoplasmic FMR1-interacting protein 2 (CYFIP2) abundance over time and partial recovery in rNLS8 mice allowed to recover for 2 weeks on dox. **E**. Differential exon usage of Unc13a at E050 correlates with a significant decrease in Unc13a expression (FKPM) and a gradual decrease in UNC13A protein abundance over time ^29^. Data represents n = 4-5 mice per group. *p <0.05 by t-test.

In the recovery cortex (2 WOD + 6 wks), *n* = 67 genes showed significant DEU, which were enriched for cell projection organisation, chemical synaptic transmission, and regulation of autophagy including TORC1 signalling (Fig. 3A). Interestingly, *n* = 48 genes were unique to the recovery cortex indicating DEU events in different genes after inhibition of neuronal hTDP-43^ΔNLS^ expression (Fig. 3B, C). We identified *n* = 19 genes with significant differential exon usage in both onset and recovery cortex, and of these *n* = 10 are neuronal and/or synaptic genes, namely *Clcn2, Fat3, Gls, L1cam, Myt1l, Nrxn1, Peg3, Siae*, *Syne1* and *Unc13a* (Fig. 3C). Interestingly however, the sites of DEU in these genes were different in onset and recovery cortex, with representative genes associated with ALS and/or FTD are illustrated in Supplementary Fig. 2, for instance Nrxn1^43^. Most of the alternatively spliced genes did not show significant changes in protein abundance or transcript levels at disease onset (Supplementary Fig. 3A). Only two downregulated genes, *Camkk2* and *Scn4b*, showed decreased protein abundance, while one upregulated gene, *Shmt1*, exhibited increased protein abundance (Supplementary Fig. 3B) comparing to the previous proteomics data^29^.

Genes with DEU included *Unc13a* and *Cyfip2*, which have previously shown alternative splicing upon TDP-43 loss of function^44^. Nevertheless, the *Stmn2* gene that has previously shown alternative splicing in response to TDP-43 pathology^17,20^ was downregulated at onset (2 WOD) but normalised at recovery, without evidence of alternative splicing. DEU of *Cyfip2* (Fig. 3D) and *Unc13a* (Fig. 3E) correlated with a decrease in the expression of these genes in rNLS8 mouse cortex, and analysis of published longitudinal proteomics data from rNLS8 mouse cortex^29^ showed a similar progressive decrease in the abundance of both proteins.

### rNLS8 mice develop hyperlocomotion, hyperactivity, and habituation deficits, concomitant with impaired motor functions, which persist even after motor recovery from advanced disease

Given the indication of hyperlocomotion during recovery from early disease in rNLS8 mice but minimal persistent molecular changes, we next sought to evaluate how extra-motor phenotypes are affected in rNLS8 mice during recovery from late stage of disease, and to determine molecular correlates of these behavioural alterations. Before characterising the extra-motor behavioural phenotypes in the rNLS8 mice, we thus extended the disease phase up to 6 WOD prior to a recovery phase of an additional 6 weeks back on dox (Fig. 4A). As expected, rNLS8 mice showed significant motor deficits at 6 WOD, and importantly, recovered motor function in inverted grid performance and grip strength and dramatically improved rotarod performance through + 6 wks back on dox (Fig. 4B), aligning with previous reports^28^.

**Figure 4.**
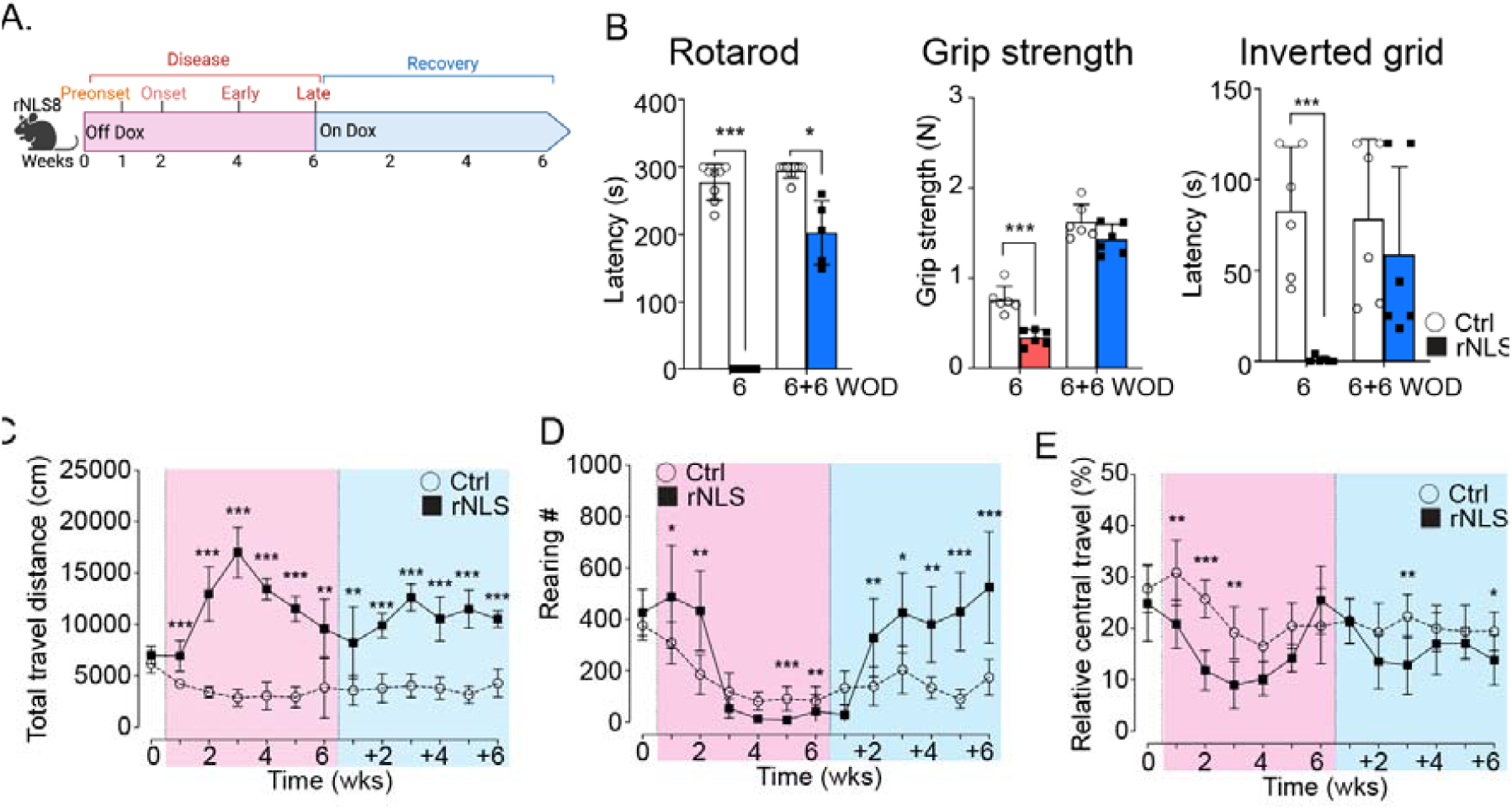
rNLS8 mice exhibit disinhibition-like phenotypes, and social and executive deficits despite impaired motor function. (**A**) The schematic of the rNLS8 mouse model. (**B**) The data of rotarod, grip strength and inverted grid tests at 6 WOD and 6 + 6 weeks back on dox. The open field analyses revealed the (**C**) total distance, (**D**) rearing number, and (**E**) relative central travel of the experimental mice that travelled in the open field arena over 30 min at baseline before the removal of dox (0 WOD), at 1-to-6 WOD and at additional 1-to-6 weeks back on dox. n ≥ 9 per group. Data as mean ± SD. * p < 0.05, ** p < 0.01 by two-way ANOVA.

In the open field test, the rNLS8 mice exhibited a significant increase in travel distance compared to the control mice at all timepoints examined, even prior to motor onset (1 WOD; Fig. 4C) and throughout recovery (+1 to +6 wks back on dox). The hyperlocomotion in the rNLS8 mice peaked at 3 WOD (Fig. 4C), in line with a recent finding^24^, then gradually decreased at later timepoints of disease correlating with increasing motor deficits in rNLS8 mice. In contrast to recovered motor function (Fig. 4B), the rNLS8 mice continued to display significant hyperlocomotion at all recovery timepoints (Fig. 4C). These results suggest the rNLS8 mice developed a disinhibition-like phenotype prior to the onset of motor deficits, which peaked at early disease stages, remained significantly increased despite severe motor deficits from 4 WOD, and then persisted during motor recovery. The rNLS8 mice also displayed dramatic increases of the travel distance in the outside areas (walls and corners) that involve exploring and walking^41^ (Supplementary Fig. 4A,B), and persistent increases of initial total travel distance (0-5 min, Supplementary Fig. 4C), suggesting hyperactivity^41^. Furthermore, time-binned analysis revealed habituation deficits^42^, at 4 WOD (5-30 min, Supplementary Fig. 4D).

rNLS8 mice also displayed a dramatic increase of rearing number in the open field test at 1 and 2 WOD (Fig. 4D) which was less evident at later timepoints when hindlimb function was impaired. Importantly, the rNLS8 mice re-displayed the increased rearing behaviour by 2 wks back on dox (Fig. 4D and Supplementary Fig. 4E), indicating persistence of this phenotype that had been masked by motor impairment in the later disease stages. We also observed significant decreases in the relative central travel (%) in the rNLS8 mice after motor onset at 3 WOD, which persisted to the recovery phase through 6 WOD +6wks, implying an anxiety-like phenotype (Fig. 4E). However, we did not observe significant space-related anxiety-like behaviours in the commonly used elevated plus maze paradigm (Supplementary Fig. 5). Nevertheless, we detected slightly decreased anxiety-like behaviour in rNLS8 mice in the light-dark transition test, with significantly decreased duration (%) in the dark chamber (Supplementary Fig. 6), in line with our previous finding in the *CAMKII*-hTDP-43^ΔNLS^ mice^33^. These data suggest the decreases of relative central travel (Fig. 4E) may be due to the dramatic increases of travel distance in the areas close to walls and corners. Together, the data revealed persistent hyperlocomotion and hyperactivity phenotypes in rNLS8 mice.

### rNLS8 mice display persistent social and executive deficits

As apathy is a prevalent feature of individuals with ALS/FTD^8,45^, we examined sociability and social memory of rNLS8 mice using three-chamber social interaction tests^45^ at 1, 2, 4 WOD and 6 WOD + 6 wks back on dox (Fig. 5A). As expected, the control mice spent more time with the novel mouse than the familiar mouse (Fig. 5B), which indicates a natural preference for novelty^45^. In contrast, rNLS8 mice displayed similar length of interaction time between novel and familiar mice, suggesting impaired social memory. Importantly, the social memory impairment persisted at 6 WOD + 6 wks back on dox (Fig. 5C), with persistent increased travel distance in the testing arena correlating with our previous findings of hyperlocomotion (Supplementary Fig. 7). Overall, the social interaction results demonstrate persistent sociability and social memory deficits in rNLS8 mice in disease and despite motor function recovery.

**Figure 5.**
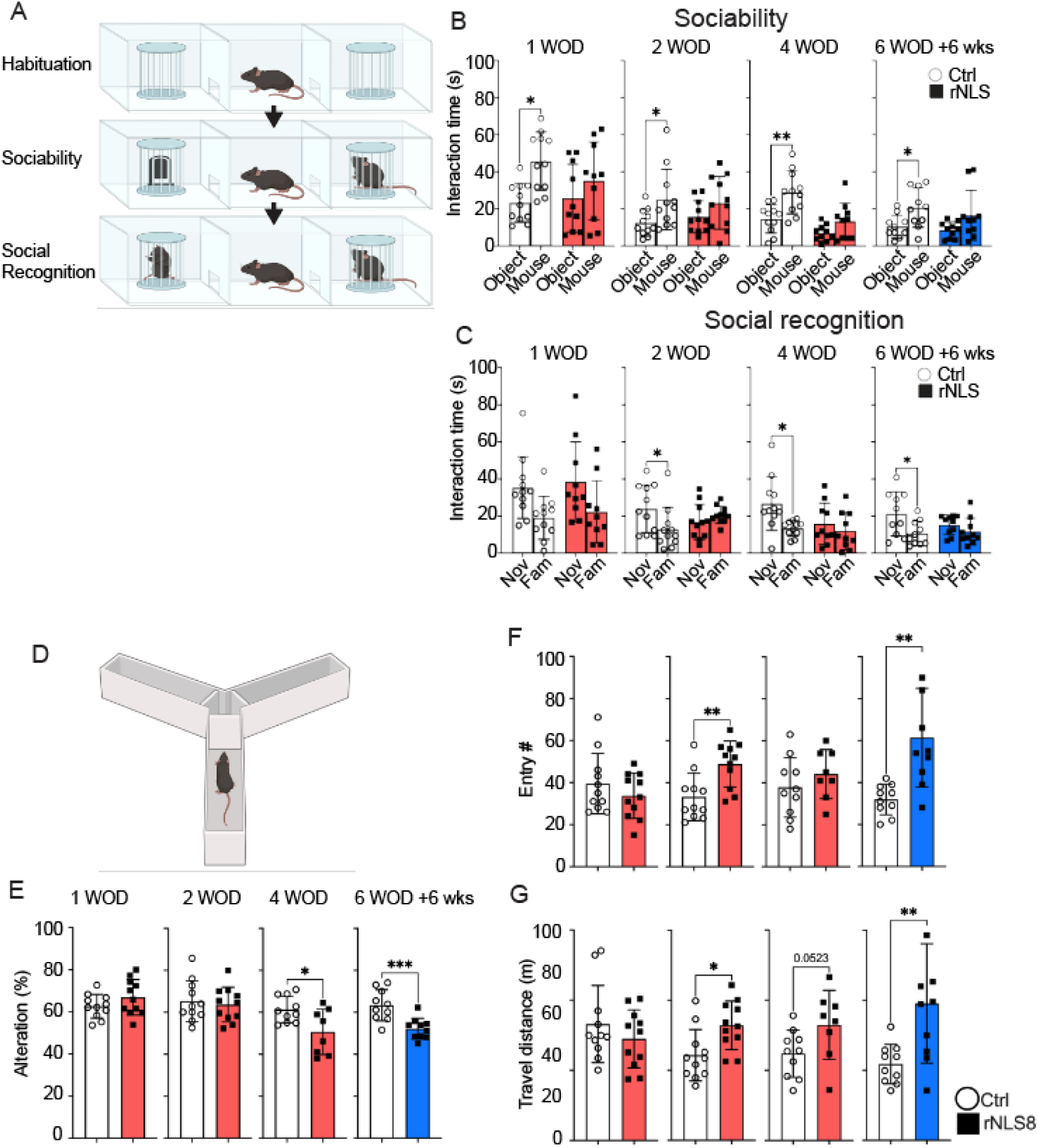
rNLS8 mice exhibit social and executive deficits despite impaired motor function. (**A**) Schematic of three-chamber social interaction tests. (**B**) Sociability session to compare the interaction time (second) that the experimental mice spent with the non-social object (Object) versus the social object (mouse) at 1, 2, 4 WOD and 6 WOD +6 weeks back on dox. (**C**) Social recognition session to assess the interaction time (second) that the experimental mice spent with the novel (Nov) social stimulus versus the familiar (Fam) social object at 1, 2, 4 WOD and +6 weeks back on dox. (**D**) Schematic of three-chamber social interaction tests**. (E)** Spontaneous alteration (%). (**F**) Total entry number and (**G**) the total travel distance (m) at 1, 2, 4 WOD and +6 weeks back on dox. n ≥ 9 per group. Data as mean ± SD. * p < 0.05, ** p < 0.01 by two-way ANOVA.

We further examined spatial working memory to assess executive function in rNLS8 mice, using the Y-maze test (Fig. 5D) as in our previous reports of *CAMKII*-hTDP-43^ΔNLS^ mice^30, 33^. rNLS8 mice displayed similar spontaneous alterations (>60%) to the control mice until disease onset, including 1 and 2 WOD (Fig. 5E). Interestingly, the rNLS8 mice started to exhibit a significant reduction of spontaneous alteration (%) comparing to controls at 4 WOD, in line with a recent report suggesting contribution of hippocampal neurodegeneration to executive deficits in the rNLS8 mice^24^. Notably, the executive deficits persisted to the recovery stage in the rNLS8 mice (Fig. 5E). Moreover, we observed the consistent increased entry number (Fig. 5F) and total travel distance (Fig. 5G) at 2 WOD and 6 WOD +6 wks back on dox, respectively, in line with the findings in the open field and social interaction tests. It is worth further investigation as to how neurodegeneration in different brain regions contributes to executive dysfunction in the rNLS8 mice and whether this is related to ALS and FTD.

### Glutamatergic signalling protein loss persists in the rNLS8 mouse cortex following motor recovery, similar to changes in human ALS/FTD

We hypothesised that synaptic mechanisms^14, 46, 47^ may contribute to the persistent behavioural changes in the rNLS8 mice, and we therefore re-analysed a previously published longitudinal proteomics dataset of the rNLS8 mouse cortex^29^. We focused on the subset of proteins that were significantly decreased in the cortex in late disease (6 WOD, *n* = 305 proteins; fold change > 1.2 and P-value < 0.05), which were enriched for biological processes associated with “chemical synaptic transmission” and “behaviour”. Approximately half of this subset of late disease decreased proteins (*n* = 117; 51%) were restored to control levels in recovery (“recovered” proteins = down at 6 WOD but no change at +2 weeks back on dox), and the other half remained decreased in recovery in rNLS8 mice (*n* = 114, 49%; “persistent” proteins = down at 6 WOD and down at +2 weeks back on dox; Fig. 6A and B; Supplementary Table 2). Interestingly, “recovered” and “persistent” subsets of proteins were both enriched for proteins associated with synapse biology (Fig. 6C), suggesting that some synaptic proteins are more “plastic” and capable of restoring to control levels than others, for example, recovered synaptic gamma-aminobutyric acid receptor subunit alpha-3 (GABRA3) that was altered in sporadic FTD-TDP cases^48^. Interestingly, persistently decreased proteins in disease and recovery were enriched for glutamatergic synapse proteins, revealed by independent gene ontology analysis approaches and a large protein-protein interaction network (Fig. 6C and D, Metascape and Ingenuity Pathway Analysis; Supplementary Table 2). Persistently decreased glutamatergic synapse proteins included syntaxin-1A (STX1A), voltage-dependent N-type calcium channel subunit alpha-1B (CACNA1B), and calcium/calmodulin-dependent protein kinase type IV (CAMK4), as well as Glutamate Ionotropic Receptor AMPA Type Subunit 2 and 3 (Gria2 and Gria3). SynGo analysis suggested alterations in both pre- and post-synaptic components (Fig. 6E).

**Figure 6.**
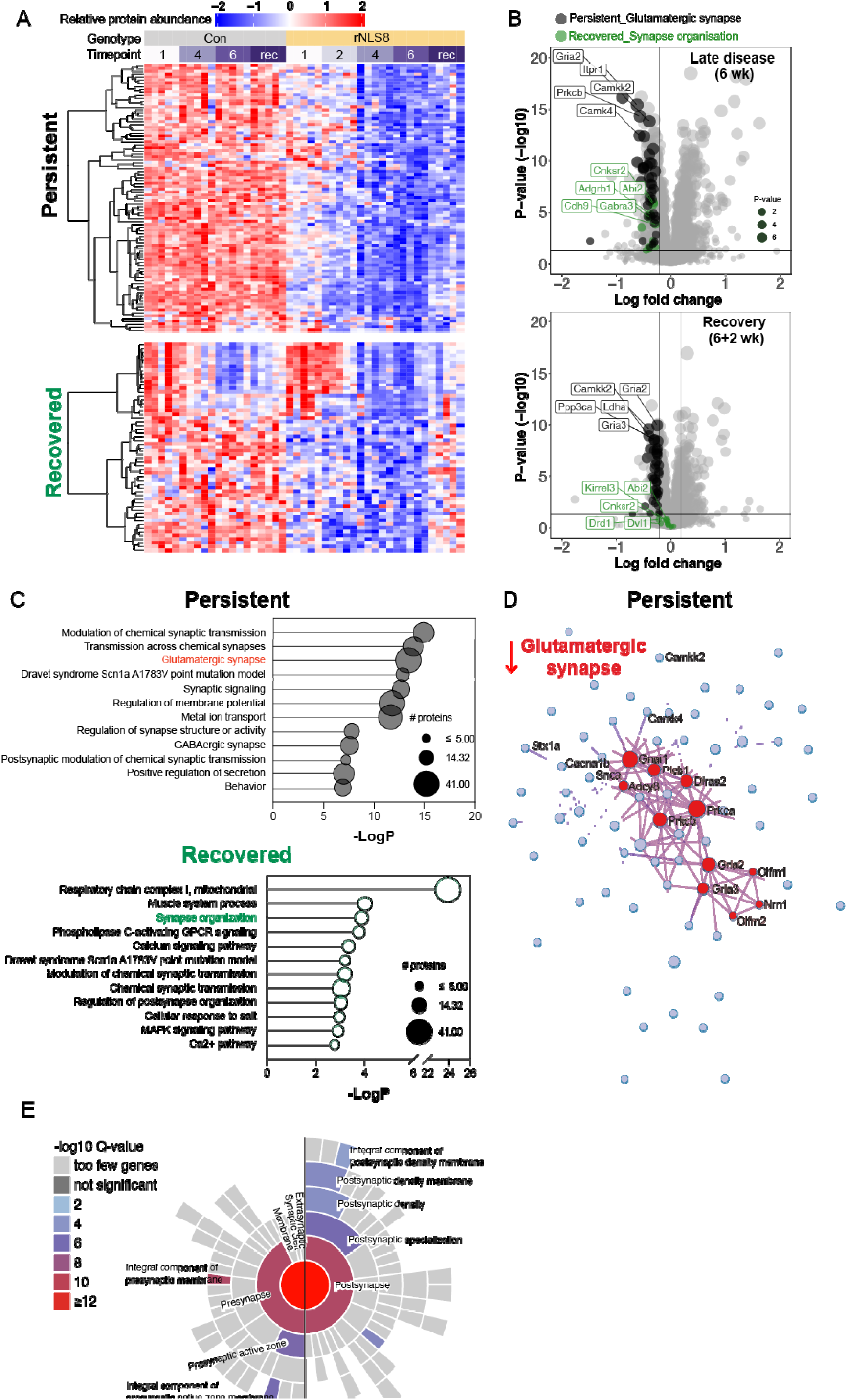
Depletion of a subset of synaptic pathway components including glutamate signalling proteins persists in rNLS8 mice despite motor recovery. (**A**) Heatmap of the relative protein abundance of quantified proteins in “persistent” and “recovered” protein subsets in rNLS8 mice at 1, 2, 4, and 6 WOD (WOD) and in recovery (6 WOD + 2 weeks on dox) compared to littermate controls (Con). The persistent subset of proteins represents those that are significantly decreased (fold change > 1.2, p-value < 0.05) in rNLS8 mice in late disease (6 WOD) and recovery (6 WOD + 2 weeks on dox). The recovered subset of proteins represents those that are significantly decreased in rNLS8 mice in late disease (6 WOD) but are not significantly different to control levels in recovery (6 WOD + 2 weeks on dox). Each column represents data from and individual mouse (n=5/group), and red = high and blue = low relative protein abundance. (**B**) Volcano plots of mean log fold change (rNLS8/Con) and p-value (-log10) from late disease (6 WOD) and recovery (6 WOD + 2 weeks on dox) timepoints. All proteins (grey), proteins belonging to the glutamatergic synapse (black) gene ontology and the synapse organisation (green) gene ontology terms are shown. (**C**) Metascape gene ontology analysis of persistent and recovered subsets of proteins. The size of the bubble indicates the number of proteins in each term. (**D**) Protein-protein interaction network of components of the glutamatergic synapse (red), which are persistently significantly decreased in disease (6 WOD) and recovery (6 WOD + 2 weeks on dox). Proteomics data^29^ were re-analysed to identify persistently decreased and recovered proteins in the rNLS8 mouse cortex. (**E**) Synaptic protein annotations of the glutamatergic synapse proteins that persistently significantly decreased in disease and recovery by SynGO^49^.

A proportion of the glutamatergic synapse proteins that were persistently decreased in rNLS8 mouse cortex in late disease and recovery were also significantly decreased in human ALS and FTD post-mortem brain tissues^38, 39^, revealed by targeted re-analysis of published proteomic and transcriptomic datasets using the webtool “TDP-map” (Fig. 7 and Supplementary Fig. 8)^27–29^. Together, these data show that a subset of disease-relevant glutamatergic synaptic proteins that are depleted in rNLS8 mouse cortex correlating with persistent extra-motor phenotypes, are also affected in human TDP-43 proteinopathies.

**Figure 7.**
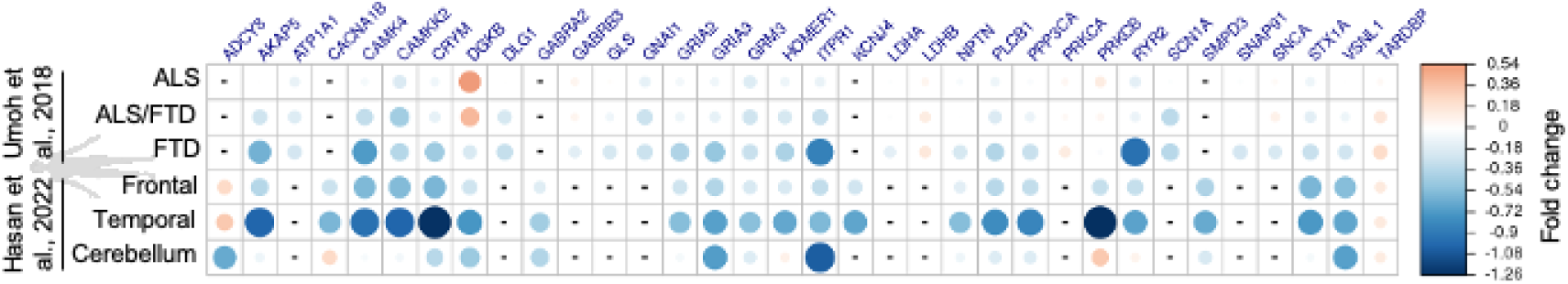
Glutamatergic synapse proteins that are persistently decreased in rNLS8 mice are also significantly decreased in human ALS and FTD post-mortem tissues. Plot of glutamatergic synapse genes/protein abundance in transcriptomic^39^ and proteomic^38^ of human post-mortem cortex was generated using the <u>TDP-map</u> webtool and datapoints are colour-scaled to illustrate the log fold change where red is increased and blue is decreased compared to controls. Circle size is dependent on the magnitude of log fold change, the larger the change the larger the circle size.

## Discussion

The growing recognition of extra-motor symptoms associated with ALS highlights their significant impact on quality of life and disease prognosis. This raises the need for understanding of the mechanisms behind these symptoms, as well as the development of animal models that accurately reflect the complexities of ALS. Here, we demonstrated that broad expression of disease-reminiscent hTDP-43^ΔNLS^ in neurons leads not only to ALS-like motor decline but also to persistent extra-motor behavioural impairments in rNLS8 mice. These persistent behavioural deficits associate with changes in splicing and levels of glutamatergic synapse proteins, suggesting potential mechanisms of relevance to disease.

### Early and persistent extra-motor behaviours in rNLS8 mice and human disease

rNLS8 mice have been well characterised as a model developing motor deficits ^26–28,50,51^, and recent studies have suggested that rNLS8 mice also develop extra-motor phenotypes including hyperactivity and anxiety deficits in early disease stages^24^. Our study reveals that rNLS8 mice display many early extra-motor behaviours that are readily detectable despite motor impairments, and further that these changes persist despite recovery of motor function upon suppression of the hTDP-43^ΔNLS^ transgene. Notably, phenotypes such as hyperlocomotion and social withdrawal emerge early, prior to the onset (1 WOD) of detectable motor impairments. Importantly, our findings reveal that extra-motor behaviours persist even during the recovery phase, when motor function, muscle innervation, and overall survival otherwise improve ^28^. This observation suggests that TDP-43 pathology rapidly inflicts lasting damage on neural circuits, possibly through ongoing synaptic dysfunction at both transcriptomic and proteomic levels^29^ (Fig. 2 and 5). While neuronal loss in vulnerable populations, such as layer V cortical neurons ^28^ and CA3 hippocampal neurons^24^, may contribute to these persistent deficits, the enduring synaptic impairments that we detected via transcriptomic analysis indicates these deficits may be caused by circuit changes rather than simply due to frank neurodegeneration.

Our results provide valuable insights that align with the extra-motor phenotypes observed in various other TDP-43 mouse models. For example, brain-specific expression of hTDP-43^ΔNLS^ driven by the *CAMKII* promoter induces hyperlocomotion and social withdrawal in mice^30, 33^. Similarly, overexpression of wildtype TDP-43 in the forebrain leads to progressive motor and memory deficits^33^. Additionally, TDP-43 mutant knock-in mice exhibit various behavioural alterations, highlighting the contributions of both gain- and loss-of-function mechanisms contributing to motor and extra-motor dysfunctions. For instance, TDP-43^G348C^ mutant mice show discernible motor and memory deficits^50^, whereas TDP-43^Q331K^ mutants present cognitive and motor impairments^51, 52^. Conversely, TDP-43^M337V^ and TDP-43^G298S^ mutants do not exhibit significant cognitive or behavioural changes ^53^, highlighting the need to carefully draw conclusions from studies of different models. Notably, the extra-motor phenotypes in rNLS8 mice align with other ALS/FTD mouse models that exhibit TDP-43 pathology, such as C9ORF72 repeat expansions (G4C2)66 mice^54^, C9BAC mice^55^, Fus^ΔNLS/+^ mice^56^, and rAAV2/8-UBQLN2 mice^57^. These findings suggest a potentially universal mechanism involving TDP-43 pathology that contributes to extra-motor dysfunction in ALS/FTD. Notably, unlike other TDP-43 models where extra-motor symptoms typically develop only after prolonged disease progression, for example TDP-43^M337V^ and TDP-43^G298S^ mice that show motor neuron degeneration only after 2.5 years^53^, the rNLS8 mouse model exhibited a rapid onset of both motor and extra-motor phenotypes.

### Synaptic mechanisms in TDP-43-associated disease

Early extra-motor changes in rNLS8 mice correlated with significant early decreases in levels of proteins associated with synaptic transmission and organisation^29^. Cortical hyperexcitability has been long proposed as a causative mechanism for ALS and FTD associated not only with TDP-43^3, 29, 58^ but other disease proteins for instance C9ORF72^47^ or FUS^56^. Notably, hyperexcitability is also a key feature of ALS^59^. In this study, we identified a hyperlocomotion and hyperactivity phenotype in the rNLS8 mice, which may be linked to early increased neuronal activity in cortical regions ^27^, preceding neurodegeneration at 3-4WOD^24, 27, 28^. Interestingly, both excitatory and inhibitory neurons displayed increased neuronal activity in layer V and layer II/III of rNLS8 mice^27^, supporting the idea that the imbalance of excitatory and inhibitory regulation contribute to disease development in the ALS^59^. A recent report^24^ showed significant hippocampal hyperexcitability in the of rNLS8 mice, implying TDP-43 pathology disrupts multiple brain regions and global circuitry. Moreover, similar hyperactivity was noted in the cortical brain section in *CAMKII*-hTDP-43^ΔNLS^ mice^60^, emphasizing the importance of exploring how TDP-43 pathology may lead to both gain- and loss-of-function that could result in hyperexcitability and consequent neurotoxicity^40^.

TDP-43, as a DNA-/RNA-binding protein, binds the transcripts of numerous synaptic genes that regulates neural circuitry vital for the cognitive and motor functions under health and disease conditions^53–55^. The role of TDP-43 in RNA metabolism contributes to disease, and targeting TDP-43 pathology can ameliorate the disease development and prolonged the life span^25^. TDP-43 loss has been implicated in the regulation of key synaptic proteins, including syntaxin-binding protein 1 (*STXBP1*), synaptotagmin-7 (*SYT7*), and *UNC13A*^19, 61^. Our data reveal that rNLS8 mice develop splicing alterations in genes enriched for axonogenesis and chemical synaptic transmission, with a subset of genes retaining DEU events during recovery phase. Notably, given that cryptic splicing of Unc13a results from loss of functional TDP-43 ^19^, correcting its splicing presents a promising therapeutic approach. A recent study also showed that rNLS8 mice develop alterations in 3’ untranslated region polyadenylation, affecting genes crucial for synaptic function and showing resemblance to human ALS/FTLD-TDP^23^.

Our study further highlights a progressive decrease in proteins related to chemical synaptic transmission in the rNLS8 cortex prior to disease onset, with particular susceptibility observed in glutamatergic synapse proteins^29^. These glutamatergic synapse proteins were particularly susceptible and significantly decreased from disease onset (1 WOD) and remained decreased in the recovery phase. This finding aligns with our previous biochemical results that cytoplasmic TDP-43 mislocalisation leads to decreased protein levels of AMPAR subunit in the rNLS8^62^ and *CAMKII*-TDP-43^ΔNLS^ mice^60^. While we have explored targeting AMPAR using riluzole, which does not effectively prevent disease phenotypes in rNLS8 mice^62^, alternative pharmacological or genetic strategies to modulate glutamatergic signalling merit investigation for therapeutic development. Targeting glutamatergic receptors could enhance synaptic activity and thereby improve cognitive function in neurodegenerative disease^63^. In addition, rNLS8 mice exhibit astrogliosis ^28, 29^, and activated astrocytes, a consistent inflammatory feature in ALS^64^, have been linked to both the pathology and motor deficits in ALS patients and TDP-43 mouse models^64–66^. The role of astrocytes in glutamate reuptake at synapses is critical for maintaining normal brain function and preventing excitotoxicity associated with disorders like ALS/FTD^64, 66^. Notably, the relationship between astrocytic modulation of synaptic function and behavioural changes, like neuronal hyperactivity and hyperlocomotion^65^, further underscores the potential impact of astrocytic dysfunction in addition to neuronal synaptic alterations on disease-relevant phenotypes in rNLS8 mice and ALS patients. Future studies could examine the underlying mechanisms using optogenetic or electrophysiological techniques.

### Limitations

The impact of TDP-43 pathology on different brain regions in addition to the cortex warrants further exploration. For example, recent studies suggest that TDP-43 pathology leads to hippocampal neuron loss in rNLS8 mice^24^, indicating that neurodegeneration of distinct neuronal subpopulations may contribute to the various behavioural phenotypes of this model. Further, consideration of differences between mouse and human pathophysiology is important. Recognizing the inherent limitations of translating findings from animal models to human conditions, we note the absence of von Economo neurons and fork cells that are key targets of degeneration in human FTD that may play a role in development of disease-relevant behavioural dysfunctions^67^, are absent in mice. Additionally, the genes containing UG-rich TDP-43 RNA binding sequences, and that are therefore splicing targets of TDP-43, show differences between mouse and human^68^. However, in this study we detected changes in levels of splicing and protein of key TDP-43-related loss of function targets, including Unc13a. Interestingly, Unc13a protein levels were decreased in the rNLS8 mice similar to decreases seen in human disease, despite the mouse genome harbouring TDP-43-binding sites in different genetic sites to human, such that in rNLS8 mice different exons in Unc13a were alternatively spliced to those that are impacted by loss of TDP-43 in human neurons. Finally, our studies of persistent behaviours and synaptic gene/protein changes extended only to six weeks after inhibition of the hTDP-43^ΔNLS^ transgene, and studies at further timepoints are warranted to examine the dynamics of neural recovery processes.

### Conclusion

Our study expands the known alterations, both molecular and behavioural, that result from accumulation of cytoplasmic TDP-43 in neurons. Notably, through a comprehensive integration of behavioural, transcriptomic, and proteomic analyses, our findings reveal that synaptic dysfunction, particularly at glutamatergic synapses in the rNLS8 mouse cortex, is associated with enduring behavioural deficits. Importantly, our results indicate that while recovery of motor function can be achieved rapidly with TDP-43 pathology clearance, the extra-motor behavioural phenotypes and their related molecular alterations may continue to persist. This work expands the behavioural approaches that may be applied in rNLS8 mice for pre-clinical assessment of potential ALS and FTD therapeutics. Our work also underscores the need for further exploration of the mechanisms driving TDP-43-mediated neurodegeneration, and highlights the importance of understanding the mechanisms of synaptic dysfunction in disease. These findings may also inform the development of therapeutic strategies to address the management of extra-motor symptoms to enhance the quality of life of individuals affected by these diseases.

## Acknowledgements

This work was supported by the Ross Maclean Fellowship, Brazil Family Program for Neurology, and a FightMND Bill Guest Mid-Career Research Fellowship to A.K.W., and MNDRA Innovator grant (IG2422) to W.L. This work was facilitated by the QBI Animal Facility and the QBI Animal Behavioural Facility at The University of Queensland. The authors thank J. Belforte for helpful comments on behavioural analysis.

## References

1. Masrori P, Van Damme P. Amyotrophic lateral sclerosis: a clinical review. Eur J Neurol 2020; 27(10): 1918–1929.

2. Bampton A, McHutchison C, Talbot K, Benatar M, Thompson AG, Turner MR. The Basis of Cognitive and Behavioral Dysfunction in Amyotrophic Lateral Sclerosis. Brain Behav 2024; 14(11): e70115.

3. Strong MJ, Abrahams S, Goldstein LH, Woolley S, McLaughlin P, Snowden J et al. Amyotrophic lateral sclerosis - frontotemporal spectrum disorder (ALS-FTSD): Revised diagnostic criteria. Amyotroph Lateral Scler Frontotemporal Degener 2017; 18(3-4): 153–174.

4. Vucic S, Ferguson TA, Cummings C, Hotchkin MT, Genge A, Glanzman R et al. Gold Coast diagnostic criteria: Implications for ALS diagnosis and clinical trial enrollment. Muscle Nerve 2021; 64(5): 532–537.

5. Crockford C, Newton J, Lonergan K, Chiwera T, Booth T, Chandran S et al. ALS-specific cognitive and behavior changes associated with advancing disease stage in ALS. Neurology 2018; 91(15): e1370–e1380.

6. Abrahams S. Neuropsychological impairment in amyotrophic lateral sclerosis-frontotemporal spectrum disorder. Nat Rev Neurol 2023; 19(11): 655–667.

7. Nguyen C, Caga J, Mahoney CJ, Kiernan MC, Huynh W. Behavioural changes predict poorer survival in amyotrophic lateral sclerosis. Brain Cogn 2021; 150: 105710.

8. Shojaie A, Rota S, Al Khleifat A, Ray Chaudhuri K, Al-Chalabi A. Non-motor symptoms in amyotrophic lateral sclerosis: lessons from Parkinson’s disease. Amyotrophic Lateral Sclerosis and Frontotemporal Degeneration 2023; 24(7-8): 562–571.

9. Beswick E, Forbes D, Hassan Z, Wong C, Newton J, Carson A et al. A systematic review of non-motor symptom evaluation in clinical trials for amyotrophic lateral sclerosis. J Neurol 2022; 269(1): 411–426.

10. Fisher EMC, Greensmith L, Malaspina A, Fratta P, Hanna MG, Schiavo G et al. Opinion: more mouse models and more translation needed for ALS. Mol Neurodegener 2023; 18(1): 30.

11. Tziortzouda P, Van Den Bosch L, Hirth F. Triad of TDP43 control in neurodegeneration: autoregulation, localization and aggregation. Nat Rev Neurosci 2021; 22(4): 197–208.

12. Colombrita C, Onesto E, Buratti E, de la Grange P, Gumina V, Baralle FE et al. From transcriptomic to protein level changes in TDP-43 and FUS loss-of-function cell models. Biochim Biophys Acta 2015; 1849(12): 1398–1410.

13. Wu LS, Cheng WC, Chen CY, Wu MC, Wang YC, Tseng YH et al. Transcriptomopathies of pre- and post-symptomatic frontotemporal dementia-like mice with TDP-43 depletion in forebrain neurons. Acta Neuropathol Commun 2019; 7(1): 50.

14. Gelon PA, Dutchak PA, Sephton CF. Synaptic dysfunction in ALS and FTD: anatomical and molecular changes provide insights into mechanisms of disease. Front Mol Neurosci 2022; 15: 1000183.

15. Ling SC. Synaptic Paths to Neurodegeneration: The Emerging Role of TDP-43 and FUS in Synaptic Functions. Neural Plast 2018; 2018: 8413496.

16. Klim JR, Williams LA, Limone F, Guerra San Juan I, Davis-Dusenbery BN, Mordes DA et al. ALS-implicated protein TDP-43 sustains levels of STMN2, a mediator of motor neuron growth and repair. Nat Neurosci 2019; 22(2): 167-179.

17. Prudencio M, Humphrey J, Pickles S, Brown AL, Hill SE, Kachergus JM et al. Truncated stathmin-2 is a marker of TDP-43 pathology in frontotemporal dementia. J Clin Invest 2020; 130(11): 6080–6092.

18. Baughn MW, Melamed Z, Lopez-Erauskin J, Beccari MS, Ling K, Zuberi A et al. Mechanism of STMN2 cryptic splice-polyadenylation and its correction for TDP-43 proteinopathies. Science 2023; 379(6637): 1140–1149.

19. Ma XR, Prudencio M, Koike Y, Vatsavayai SC, Kim G, Harbinski F et al. TDP-43 represses cryptic exon inclusion in the FTD-ALS gene UNC13A. Nature 2022; 603(7899): 124–130.

20. Agra Almeida Quadros AR, Li Z, Wang X, Ndayambaje IS, Aryal S, Ramesh N et al. Cryptic splicing of stathmin-2 and UNC13A mRNAs is a pathological hallmark of TDP-43-associated Alzheimer’s disease. Acta Neuropathol 2024; 147(1): 9.

21. Bademosi AT, Walker AK. Cryptic inclusions UNCover losses driving neurodegeneration. Trends Genet 2022; 38(9): 889–891.

22. Brown AL, Wilkins OG, Keuss MJ, Hill SE, Zanovello M, Lee WC et al. TDP-43 loss and ALS-risk SNPs drive mis-splicing and depletion of UNC13A. Nature 2022; 603(7899): 131–137.

23. Eck RJ, Valdmanis PN, Liachko NF, Kraemer BC. Alternative 3’ UTR polyadenylation is disrupted in the rNLS8 mouse model of ALS/FTLD. Mol Brain 2025; 18(1): 1.

24. Rodemer W, Ra I, Jia E, Gujral J, Zhang B, Hoxha K et al. Hyperexcitability precedes CA3 hippocampal neurodegeneration in a dox-regulatable TDP-43 mouse model of ALS-FTD. bioRxiv 2024.

25. Droppelmann CA, Campos-Melo D, Noches V, McLellan C, Szabla R, Lyons TA et al. Mitigation of TDP-43 toxic phenotype by an RGNEF fragment in amyotrophic lateral sclerosis models. Brain 2024; 147(6): 2053–2068.

26. Luan W, Wright AL, Brown-Wright H, Le S, San Gil R, Madrid San Martin L et al. Early activation of cellular stress and death pathways caused by cytoplasmic TDP-43 in the rNLS8 mouse model of ALS and FTD. Mol Psychiatry 2023; 28(6): 2445–2461.

27. Xie M, Miller AS, Pallegar PN, Umpierre A, Liang Y, Wang N et al. Rod-shaped microglia interact with neuronal dendrites to regulate cortical excitability in TDP-43 related neurodegeneration. bioRxiv 2024.

28. Walker AK, Spiller KJ, Ge G, Zheng A, Xu Y, Zhou M et al. Functional recovery in new mouse models of ALS/FTLD after clearance of pathological cytoplasmic TDP-43. Acta Neuropathol 2015; 130(5): 643–660.

29. San Gil R, Pascovici D, Venturato J, Brown-Wright H, Mehta P, Madrid San Martin L et al. A transient protein folding response targets aggregation in the early phase of TDP-43-mediated neurodegeneration. Nat Commun 2024; 15(1): 1508.

30. Alfieri JA, Pino NS, Igaz LM. Reversible behavioral phenotypes in a conditional mouse model of TDP-43 proteinopathies. J Neurosci 2014; 34(46): 15244–15259.

31. Zamani A, Walker AK, Rollo B, Ayers KL, Farah R, O’Brien TJ et al. Impaired glymphatic function in the early stages of disease in a TDP-43 mouse model of amyotrophic lateral sclerosis. Transl Neurodegener 2022; 11(1): 17.

32. Yang M, Silverman JL, Crawley JN. Automated three-chambered social approach task for mice. Curr Protoc Neurosci 2011; Chapter 8: Unit 8 26.

33. Alfieri JA, Silva PR, Igaz LM. Early Cognitive/Social Deficits and Late Motor Phenotype in Conditional Wild-Type TDP-43 Transgenic Mice. Front Aging Neurosci 2016; 8: 310.

34. Anders S, Reyes A, Huber W. Detecting differential usage of exons from RNA-seq data. Genome Res 2012; 22(10): 2008–2017.

35. Gu Z, Eils R, Schlesner M. Complex heatmaps reveal patterns and correlations in multidimensional genomic data. Bioinformatics 2016; 32(18): 2847–2849.

36. Zhou Y, Zhou B, Pache L, Chang M, Khodabakhshi AH, Tanaseichuk O et al. Metascape provides a biologist-oriented resource for the analysis of systems-level datasets. Nat Commun 2019; 10(1): 1523.

37. Krämer A, Green J, Pollard J, Jr, Tugendreich S. Causal analysis approaches in Ingenuity Pathway Analysis. Bioinformatics 2013; 30(4): 523–530.

38. Umoh ME, Dammer EB, Dai J, Duong DM, Lah JJ, Levey AI et al. A proteomic network approach across the ALS-FTD disease spectrum resolves clinical phenotypes and genetic vulnerability in human brain. EMBO Mol Med 2018; 10(1): 48–62.

39. Hasan R, Humphrey J, Bettencourt C, Newcombe J, Consortium NA, Lashley T et al. Transcriptomic analysis of frontotemporal lobar degeneration with TDP-43 pathology reveals cellular alterations across multiple brain regions. Acta Neuropathol 2022; 143(3): 383–401.

40. Odierna GL, Vucic S, Dyer M, Dickson T, Woodhouse A, Blizzard C. How do we get from hyperexcitability to excitotoxicity in amyotrophic lateral sclerosis? Brain 2024; 147(5): 1610–1621.

41. Choleris E, Thomas AW, Kavaliers M, Prato FS. A detailed ethological analysis of the mouse open field test: effects of diazepam, chlordiazepoxide and an extremely low frequency pulsed magnetic field. Neurosci Biobehav Rev 2001; 25(3): 235–260.

42. Konsolaki E, Skaliora I. Motor vs. cognitive elements of apparent “hyperlocomotion”: a conceptual and experimental clarification. Proc Natl Acad Sci U S A 2015; 112(1): E3–4.

43. Fernando MB, Fan Y, Zhang Y, Tokolyi A, Murphy AN, Kammourh S et al. Phenotypic complexities of rare heterozygous neurexin-1 deletions. bioRxiv 2024.

44. Ma XR, Prudencio M, Koike Y, Vatsavayai SC, Kim G, Harbinski F et al. TDP-43 represses cryptic exon inclusion in the FTD–ALS gene UNC13A. Nature 2022; 603(7899): 124-130.

45. Jabarin R, Netser S, Wagner S. Beyond the three-chamber test: toward a multimodal and objective assessment of social behavior in rodents. Mol Autism 2022; 13(1): 41.

46. Clayton EL, Huggon L, Cousin MA, Mizielinska S. Synaptopathy: presynaptic convergence in frontotemporal dementia and amyotrophic lateral sclerosis. Brain 2024; 147(7): 2289–2307.

47. Starr A, Sattler R. Synaptic dysfunction and altered excitability in C9ORF72 ALS/FTD. Brain Res 2018; 1693(Pt A): 98–108.

48. Andres-Benito P, Gelpi E, Povedano M, Santpere G, Ferrer I. Gene Expression Profile in Frontal Cortex in Sporadic Frontotemporal Lobar Degeneration-TDP. J Neuropathol Exp Neurol 2018; 77(7): 608–627.

49. Koopmans F, van Nierop P, Andres-Alonso M, Byrnes A, Cijsouw T, Coba MP et al. SynGO: An Evidence-Based, Expert-Curated Knowledge Base for the Synapse. Neuron 2019; 103(2): 217–234 e214.

50. Swarup V, Phaneuf D, Bareil C, Robertson J, Rouleau GA, Kriz J et al. Pathological hallmarks of amyotrophic lateral sclerosis/frontotemporal lobar degeneration in transgenic mice produced with TDP-43 genomic fragments. Brain 2011; 134(Pt 9): 2610–2626.

51. White MA, Kim E, Duffy A, Adalbert R, Phillips BU, Peters OM et al. TDP-43 gains function due to perturbed autoregulation in a Tardbp knock-in mouse model of ALS-FTD. Nat Neurosci 2018; 21(4): 552–563.

52. Wong P, Ho WY, Yen YC, Sanford E, Ling SC. The vulnerability of motor and frontal cortex-dependent behaviors in mice expressing ALS-linked mutation in TDP-43. Neurobiol Aging 2020; 92: 43–60.

53. Ebstein SY, Yagudayeva I, Shneider NA. Mutant TDP-43 Causes Early-Stage Dose-Dependent Motor Neuron Degeneration in a TARDBP Knockin Mouse Model of ALS. Cell Rep 2019; 26(2): 364–373 e364.

54. Chew J, Gendron TF, Prudencio M, Sasaguri H, Zhang YJ, Castanedes-Casey M et al. Neurodegeneration. C9ORF72 repeat expansions in mice cause TDP-43 pathology, neuronal loss, and behavioral deficits. Science 2015; 348(6239): 1151–1154.

55. Kahriman A, Bouley J, Tuncali I, Dogan EO, Pereira M, Luu T et al. Repeated mild traumatic brain injury triggers pathology in asymptomatic C9ORF72 transgenic mice. Brain 2023; 146(12): 5139–5152.

56. Scekic-Zahirovic J, Sanjuan-Ruiz I, Kan V, Megat S, De Rossi P, Dieterle S et al. Cytoplasmic FUS triggers early behavioral alterations linked to cortical neuronal hyperactivity and inhibitory synaptic defects. Nat Commun 2021; 12(1): 3028.

57. Ceballos-Diaz C, Rosario AM, Park HJ, Chakrabarty P, Sacino A, Cruz PE et al. Viral expression of ALS-linked ubiquilin-2 mutants causes inclusion pathology and behavioral deficits in mice. Mol Neurodegener 2015; 10: 25.

58. Akcimen F, Lopez ER, Landers JE, Nath A, Chio A, Chia R et al. Amyotrophic lateral sclerosis: translating genetic discoveries into therapies. Nat Rev Genet 2023; 24(9): 642–658.

59. Menon P, Geevasinga N, van den Bos M, Yiannikas C, Kiernan MC, Vucic S. Cortical hyperexcitability and disease spread in amyotrophic lateral sclerosis. Eur J Neurol 2017; 24(6): 816–824.

60. Dyer MS, Reale LA, Lewis KE, Walker AK, Dickson TC, Woodhouse A et al. Mislocalisation of TDP-43 to the cytoplasm causes cortical hyperexcitability and reduced excitatory neurotransmission in the motor cortex. J Neurochem 2021; 157(4): 1300–1315.

61. Estades Ayuso V, Pickles S, Todd T, Yue M, Jansen-West K, Song Y et al. TDP-43-regulated cryptic RNAs accumulate in Alzheimer’s disease brains. Mol Neurodegener 2023; 18(1): 57.

62. Wright AL, Della Gatta PA, Le S, Berning BA, Mehta P, Jacobs KR et al. Riluzole does not ameliorate disease caused by cytoplasmic TDP-43 in a mouse model of amyotrophic lateral sclerosis. Eur J Neurosci 2021; 54(6): 6237–6255.

63. Hanson JE, Ma K, Elstrott J, Weber M, Saillet S, Khan AS et al. GluN2A NMDA Receptor Enhancement Improves Brain Oscillations, Synchrony, and Cognitive Functions in Dravet Syndrome and Alzheimer’s Disease Models. Cell Rep 2020; 30(2): 381–396 e384.

64. Ziff OJ, Clarke BE, Taha DM, Crerar H, Luscombe NM, Patani R. Meta-analysis of human and mouse ALS astrocytes reveals multi-omic signatures of inflammatory reactive states. Genome Res 2022; 32(1): 71–84.

65. Nagai J, Rajbhandari AK, Gangwani MR, Hachisuka A, Coppola G, Masmanidis SC et al. Hyperactivity with Disrupted Attention by Activation of an Astrocyte Synaptogenic Cue. Cell 2019; 177(5): 1280–1292 e1220.

66. Yamanaka K, Komine O. The multi-dimensional roles of astrocytes in ALS. Neurosci Res 2018; 126: 31–38.

67. Seeley WW, Carlin DA, Allman JM, Macedo MN, Bush C, Miller BL et al. Early frontotemporal dementia targets neurons unique to apes and humans. Ann Neurol 2006; 60(6): 660–667.

68. Jeong YH, Ling JP, Lin SZ, Donde AN, Braunstein KE, Majounie E et al. Tdp-43 cryptic exons are highly variable between cell types. Mol Neurodegener 2017; 12(1): 13.

