## Supplementary Material for "Synaptic changes contribute to persistent extra-motor behaviour deficits in the rNLS8 TDP-43 mouse model of amyotrophic lateral sclerosis"

**Supplementary Methods and Materials**

**Elevated plus maze test**

Anxiety-like behaviour was assessed as described (Alfieri JA 2016) using an elevated plus maze consisting of two open arms (30 cm × 6 cm × 0.3 cm) and two closed arms (30 cm × 6 cm × 15 cm) with opaque walls. The apparatus was elevated 40 cm above the floor, and the duration of the test was 5 min. The maze was placed in the centre of a homogenously illuminated room (2 m × 1.8 m; 100 lux across arms). At the beginning of the test, mice were placed in the central square facing the open arm opposite to the investigator. Number of open arm entries, percentage of time in open arms, total arm entries and total distance travelled was measured.

**Light-dark transition test**

The open field chamber as mentioned above was divided equally into the light side that was brightly illuminated by white diodes (390 lux), and the dark side that was is illuminated at 2 lux. Mice are placed into the arena facing the dark side and the door is opened automatically 3 seconds after the mouse is detected by the infrared camera. The door is used so that the mice do not enter the light chamber immediately after the release with their motivation to escape from experimenter, since the latency to enter the light chamber may serve as an index of anxiety-like behaviour. Animals were placed toward the dark chamber and allowed to freely explore for 30 min. The distance travelled in each chamber, the total number of transitions, the time spent in the each chamber, and the latency to enter the light chamber recorded by EthoVision XT software.

**Supplementary Figures**

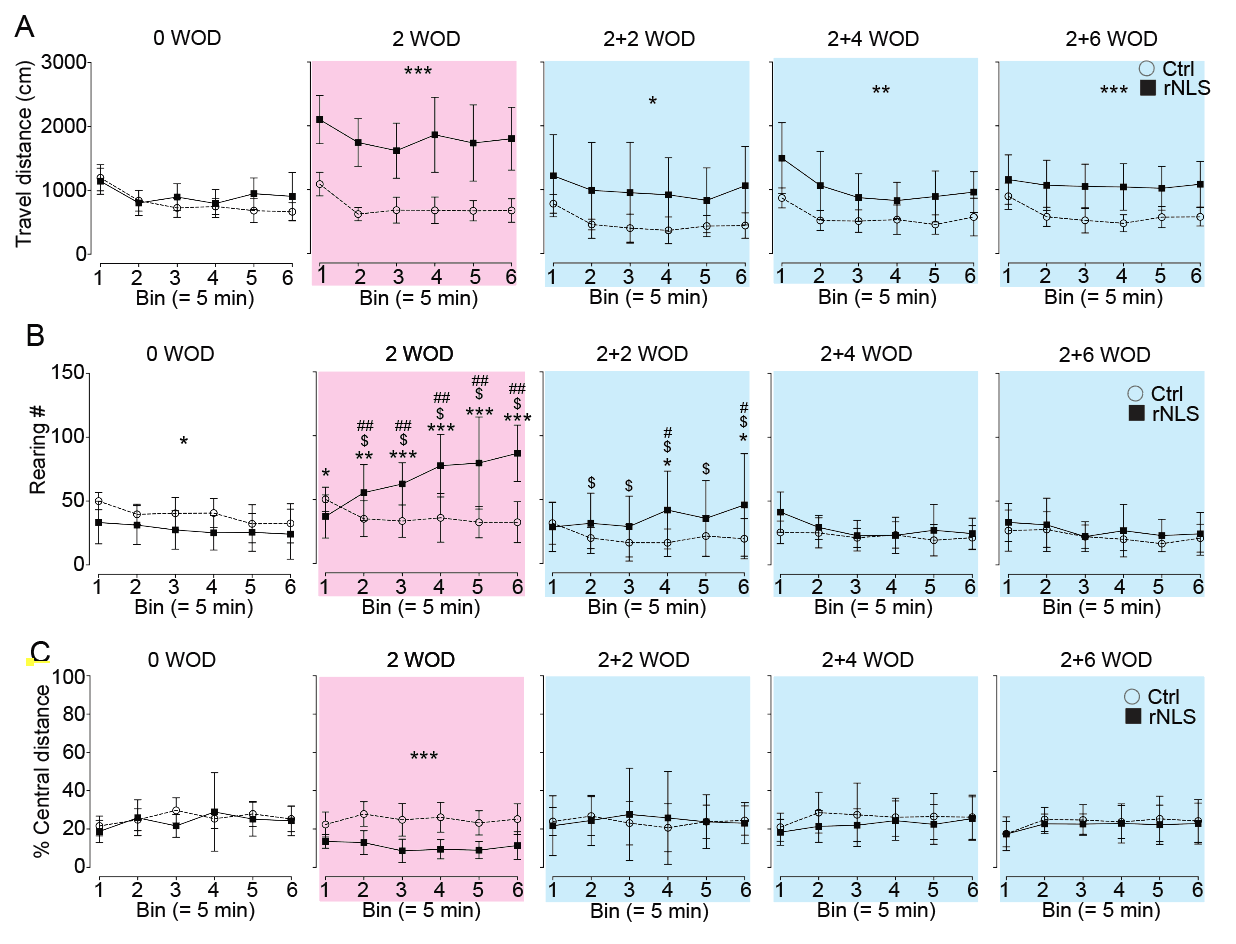
***Supplementary Figure 1.*** *Open field analysis of rNLS8 mice over weeks.* ***A****. The travel distance (cm) per bin ( = 5 min) that the experimental mice travelled in the open field arena at 0 WOD, 2 WOD, 2 WOD +2 wks back on dox , 2 WOD +4 wks back on dox, and 2 WOD + 6 wks back on dox.* ***B****. The number of rearing of experimental mice in the open field arena per bin ( = 5 min) at 0 WOD, 2 WOD, 2 WOD +2 wks back on Dox, 2 WOD +4 wks back on Dox , and 2 WOD + 6 wks back on Dox.* ***C****. The relative central distance (%) that the experimental mice travelled in the open field arena per bin ( = 5 min) at 0, 2, 2 +2, 2 +4, and 2+ 6. Data as mean ± SD. N = 11. * as p < 0.05, ** p < 0.01, *** p < 0.001 by repeated one-way ANOVA.*

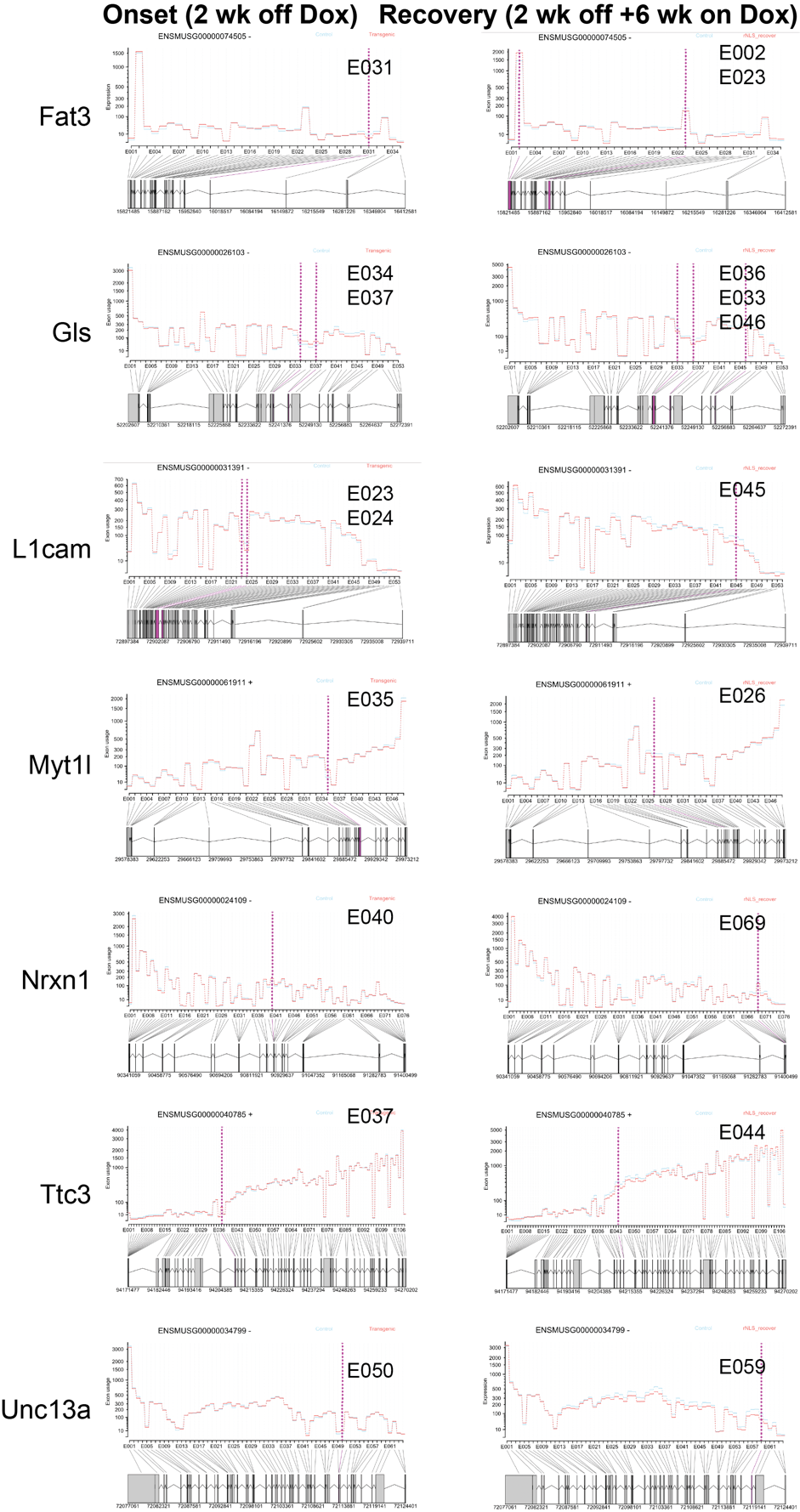

***Supplementary Figure 2.*** ***Sites of differential exon usage in alternatively splicing in representative neuronal genes associated with ALS/FTD, which were common to disease onset and recovery cortex in rNLS8 mice.*** *Left panels are onset (2 WOD) and right panels are recovery (2 WOD + 6 weeks back on Dox). Splicing plots showing exon usage at each exon feature.*

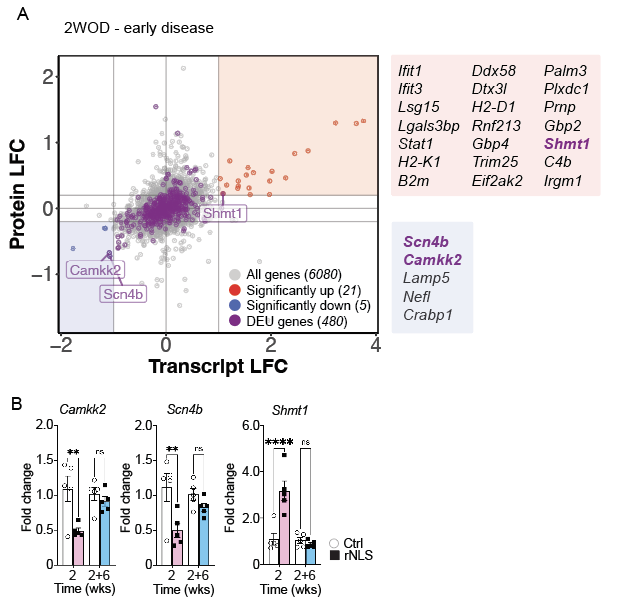

***Supplemental Figure 3. Most alternatively spliced genes are not significantly altered at the protein or transcript level at disease onset in rNLS8 mice.*** ***A****. Scatter plot correlating protein and transcript abundance (log fold change; LFC) in the cortex. All genes with corresponding protein abundance data are grey and genes with significant differential exon usage (DEU) are purple. Genes with significantly increased or decreased protein^1^ and transcripts by transcriptomic data (from this study) are red and blue, respectively, and are provided as lists ranked by their protein abundance. Scn4b and Camkk2 are genes that show DEU and have significantly decreased protein and transcript abundance. Shmt1 shows DEU and has significantly increased protein and transcript abundance. B. QPCR analyses confirmed the significant downregulation of Camkk2 and Scn4b and upregulation of Shmt1 at onset (2 WOD), which in each case was followede normalisation of levels at recovery (2 WOD + 6 weeks back on Dox). N = 5. Mean ± SEM. ** as p< 0.01 and **** as p < 0.0001 by t-test.*

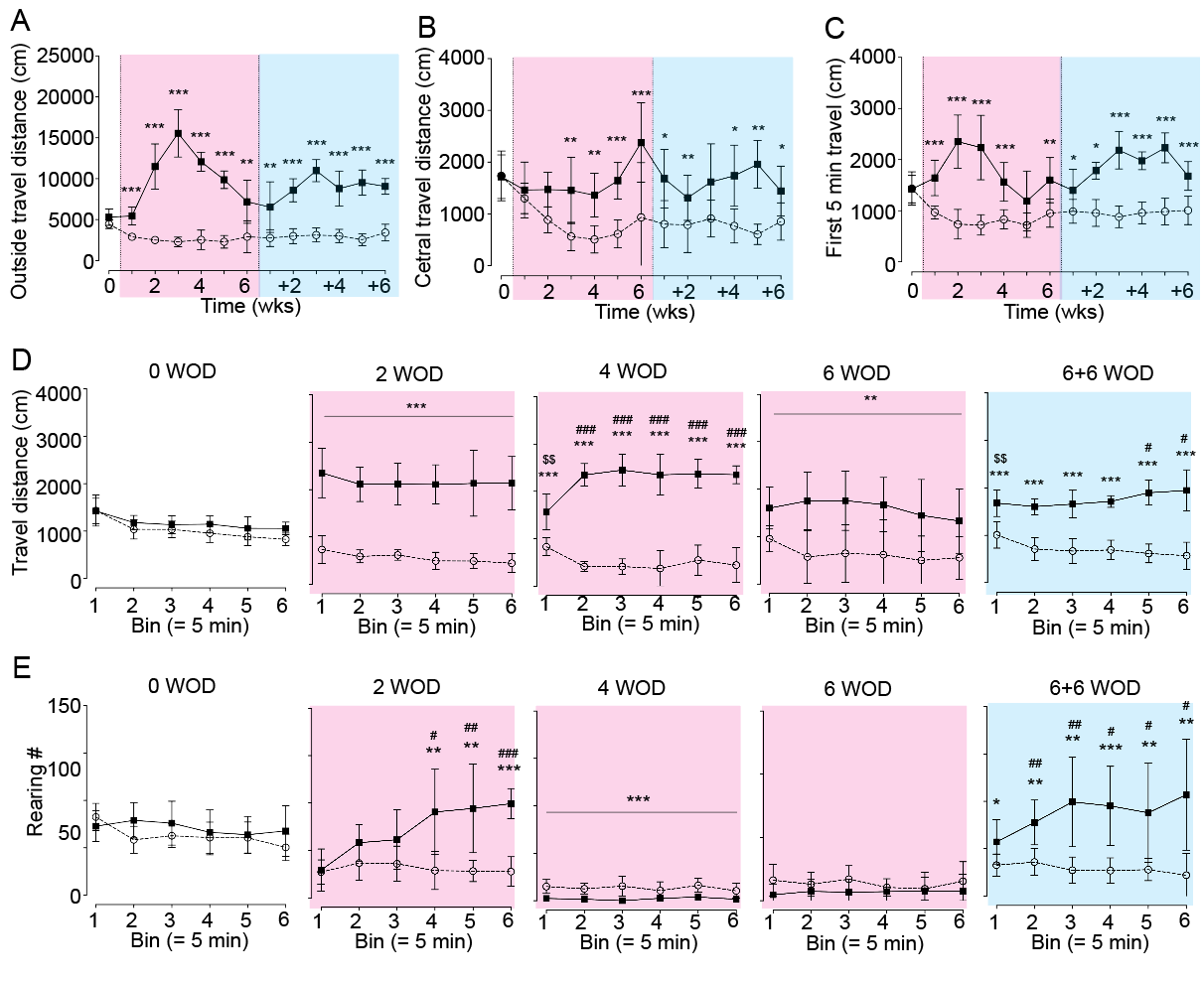
***Supplementary Figure 4.*** *rNLS8 mice display hyperlocomotion, hyperactivity phenotypes. (A) increase outside travel distance and (B) inside travel distance.* ***(C) Total travel distance in*** *the initial five minutes of the open field test during disease. The temporal profile of travel distance (cm,* ***D****) and rearing number (****E****) per bin ( = 5 min) that the experimental mice travelled in the open field arena at timepoints (****M****). Control (n = 8), rNLS8 (n =6). Mean ± SD. * as p < 0.05, ** p< 0.01, *** p < 0.001, **** p < 0.0001 between control and rNLS8 groups by repeated t-test.* $ *as p < 0.05,* $$ *p< 0.01 between first bin (=5 min) and other bins within the control group revealed by Post-hoc tests;* # *as p < 0.05,* ## *p< 0.01, #*## *p< 0.01 between first bin (=5 min) and other bins within the rNLS8 group revealed by Post-hoc tests.*

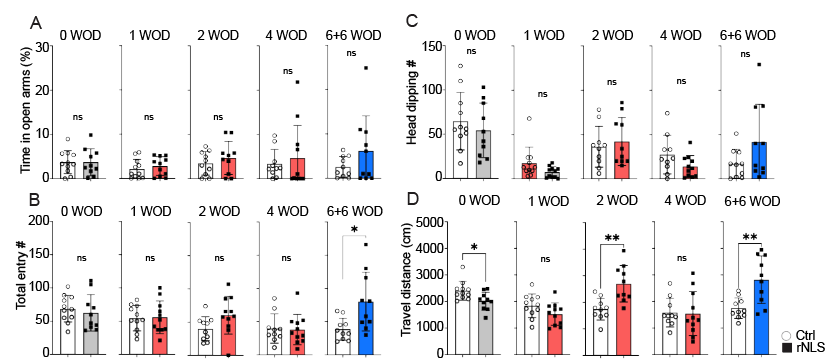

***Supplementary Figure 5.*** *rNLS8 mice exhibit no significant anxiety-like behaviours in the elevated plus maze test. (****A****) relative* *time spent (%) in the open arms. (****B)*** *Head dipping number. rNLS8 mice show slightly increased total entry number (****C)*** *at recovery phase, and increased travel distance (cm) at disease onset and recovery phase (****D)****. n ≥ 9. Data as mean ± SD. * as p < 0.05, ** p < 0.01 by t-test.*

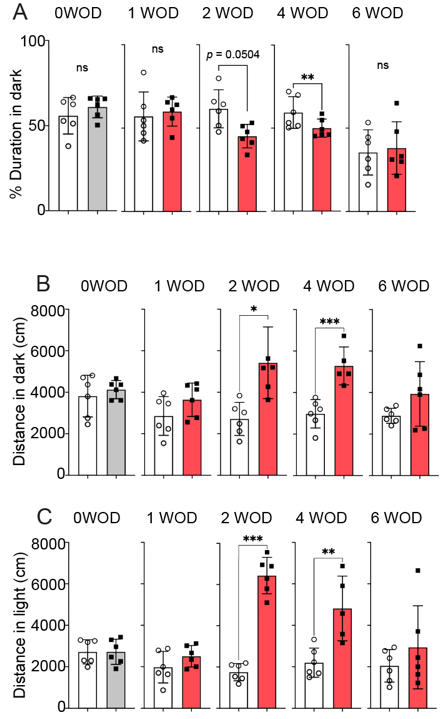

***Supplementary Figure 6. rNLS8 mice exhibit signs of reduced anxiety-like behaviour related to light. (A)*** *The relative duration (%), (B) The total travel distance (cm) in the testing chamber. (****C****) The travel distance in the dark area. (****D****) The light are in the open field arena divided equally by a light and dark area over 30 min at baseline before the removal of Dox (0 week off Dox), at 2, 4 and 6 weeks off Dox (WOD). Data as mean ± SD. N = 6. * as p < 0.05, * p < 0.01, *** p < 0.001 by t-test.*

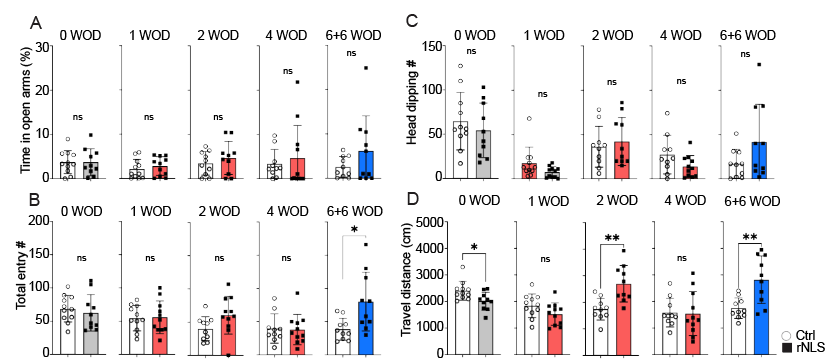

***Supplementary Figure 7. Duration and locomotor activities in social interaction testing chambers during disease and recovery phases.*** *The time (second) that the experimental mice spent in the chamber holding the non-social object (Object) versus the one holding the social object (mouse)* *at 1, 2, 4 WOD and 6 WOD +6 weeks back on Dox in the sociability session (****A****) and the social recognition session (****B****).n = 11. * as p < 0.05, ** p < 0.01 by two-way ANOVA. The locomotion was measured by the travel distance (cm) that the experimental mice spent in the chamber holding the non-social object (Object) versus the one holding the social object (mouse)* *at 1, 2, 4 WOD and 6 WOD +6 weeks back on Dox in the sociability session (****C****) and the social recognition session (****D****). N = 11. Data as mean ± SD. * as p < 0.05, ** p < 0.01, *** p < 0.01 by t-test.*

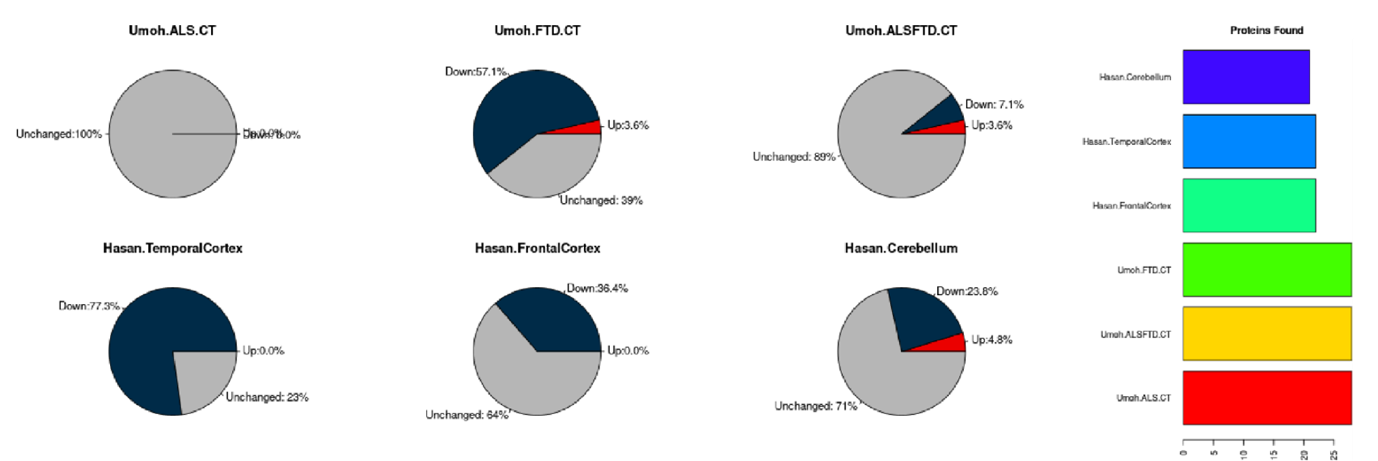

***Supplementary Figure 8****. Glutamatergic synapse proteins that are persistently decreased in rNLS8 late disease and recovery phase are also significantly decreased in the proteomics^2^ and transcriptomics^3^ datasets from human post-mortem tissue.*

**Supplementary Material References**

1.	San Gil R, Pascovici D, Venturato J, Brown-Wright H, Mehta P, Madrid San Martin L *et al.* A transient protein folding response targets aggregation in the early phase of TDP-43-mediated neurodegeneration. *Nat Commun* 2024; **15**(1)**:** 1508.

2.	Umoh ME, Dammer EB, Dai J, Duong DM, Lah JJ, Levey AI *et al.* A proteomic network approach across the ALS-FTD disease spectrum resolves clinical phenotypes and genetic vulnerability in human brain. *EMBO Mol Med* 2018; **10**(1)**:** 48-62.

3.	Hasan R, Humphrey J, Bettencourt C, Newcombe J, Consortium NA, Lashley T *et al.* Transcriptomic analysis of frontotemporal lobar degeneration with TDP-43 pathology reveals cellular alterations across multiple brain regions. *Acta Neuropathol* 2022; **143**(3)**:** 383-401.
